# Small molecule correctors divert CFTR-F508del from ERAD by stabilizing sequential folding states

**DOI:** 10.1101/2023.09.15.556420

**Authors:** Celeste Riepe, Magda Wąchalska, Kirandeep K. Deol, Anais K. Amaya, Matthew H. Porteus, James A. Olzmann, Ron R. Kopito

## Abstract

Over 80% of people with cystic fibrosis (CF) carry the F508del mutation in the cystic fibrosis transmembrane conductance regulator (CFTR), a chloride ion channel at the apical plasma membrane (PM) of epithelial cells. F508del impairs CFTR folding causing it to be destroyed by endoplasmic reticulum associated degradation (ERAD). Small molecule correctors, which act as pharmacological chaperones to divert CFTR-F508del from ERAD, are the primary strategy for treating CF, yet corrector development continues with only a rudimentary understanding of how ERAD targets CFTR-F508del. We conducted genome-wide CRISPR/Cas9 knockout screens to systematically identify the molecular machinery that underlies CFTR-F508del ERAD. Although the ER-resident ubiquitin ligase, RNF5 was the top E3 hit, knocking out *RNF5* only modestly reduced CFTR-F508del degradation. Sublibrary screens in an *RNF5* knockout background identified RNF185 as a redundant ligase, demonstrating that CFTR-F508del ERAD is highly buffered. Gene-drug interaction experiments demonstrated that correctors tezacaftor (VX-661) and elexacaftor (VX-445) stabilize sequential, RNF5-resistant folding states. We propose that binding of correctors to nascent CFTR-F508del alters its folding landscape by stabilizing folding states that are not substrates for RNF5-mediated ubiquitylation.

**SIGNIFICANCE STATEMENT:**

- Clinically effective small molecule cystic fibrosis (CF) correctors divert mutant CFTR molecules from ER-associated degradation (ERAD). However, the mechanisms underlying CFTR ERAD are not well-understood.
- The authors used CRISPR knockout screens to identify ERAD machinery targeting CFTR-F508del and found that the pathway is highly buffered, with RNF185 serving as a redundant ubiquitin ligase for RNF5. Gene-drug interaction experiments demonstrated that correctors act synergistically by stabilizing sequential RNF5-resistant folding states.
- Inhibiting proteostasis machinery is a complementary approach for enhancing current CF corrector therapies.

## INTRODUCION

Cystic fibrosis (CF) is caused by loss-of-function mutations to the gene encoding the cystic fibrosis transmembrane conductance regulator (CFTR), a chloride ion channel expressed at the apical plasma membrane (PM) of exocrine epithelial cells (Quinton, 1983; Riordan *et al*., 1989). CFTR is a large, polytopic ABC transporter composed of two transmembrane domains (TMD1 and TMD2), cytosolic regulatory (R) and ATP-binding (NBD1 and NBD2) domains (Riordan *et al*., 1989) and an interfacial lasso motif (Zhang and Chen, 2016). The most common disease mutation, F508del, found in ∼ 80% of people with CF (pwCF) (Bobadilla *et al*., 2002; De Boeck *et al*., 2014; Lopes-Pacheco, 2019; Patient Registry), impairs the folding of NBD1, leading nascent CFTR to be triaged by ER-associated degradation (ERAD) and degraded by the ubiquitin proteasome system (UPS) before it can leave the ER (Jensen *et al*., 1995; Ward *et al*., 1995). The discovery that CFTR-F508del can traffic to the PM at reduced temperatures (Denning *et al*., 1992) or in the presence of chemical chaperones like glycerol (Sato *et al*., 1996) contributed to the recognition that protein conformational disorders can be treated with small molecules that promote near-native folding and function (Bernier *et al*., 2004; Aymami *et al*., 2013; Muntau *et al*., 2014; Grasso *et al*., 2023). Consequently, high-throughput screens (HTS) of chemical libraries have been employed to identify small molecule “correctors” that rescue CFTR-F508del folding, trafficking, and function (Pedemonte *et al*., 2005; Van Goor *et al*., 2006, 2011). To date, three correctors, lumacaftor (VX-809), tezacaftor (VX-661), and elexacaftor (VX-445) have been approved for clinical use (Wainwright *et al*., 2015; Heijerman *et al*., 2019; Middleton *et al*., 2019).

Correctors can promote CFTR-F508del trafficking to the PM by either acting as “pharmacological chaperones” or “proteostasis modulators” (Lukacs and Verkman, 2012). Pharmacological chaperones directly bind to CFTR-F508del and stabilize a more native conformation while proteostasis modulators indirectly increase the efficiency of CFTR-F508del folding by influencing the protein quality control (PQC) machinery that mediates folding and assembly of nascent CFTR-F508del. Currently, all clinically-approved correctors are pharmacological chaperones, with the most potent CF therapeutic, Trikafta, relying on the synergistic interaction of elexacaftor and tezacaftor in combination with the CFTR channel gating “potentiator,” ivacaftor (Heijerman *et al*., 2019; Middleton *et al*., 2019). The approval of Trikafta in 2019 has revolutionized CF therapeutics, dramatically reducing the frequency of pulmonary exacerbations, life-threatening bacterial infections, and lung transplants for pwCF (2022 CFF Patient Registry Handout).

Individually, tezacaftor and elexacaftor produce a modest increase in CFTR-F508del function, yet together, they can restore CFTR-F508del activity to ∼50-70% WT levels (Veit *et al*., 2020; Capurro *et al*., 2021). Structural studies of CFTR and CFTR-F508del in complex with correctors reveal that tezacaftor and its congener lumacaftor bind to a hydrophobic pocket in TMD1, stabilizing four transmembrane helices that are otherwise intrinsically unstable (Fiedorczuk and Chen, 2022a) while elexacaftor stabilizes the interdomain interface of two transmembrane helices in TMD2 and the lasso motif, promoting native intramolecular dimerization of the two NBDs (Fiedorczuk and Chen, 2022b). CFTR folding begins co-translationally (Kleizen *et al*., 2021) and proceeds in at least two distinct stages (Im *et al*., 2023). Tezacaftor is likely to bind during the first, cotranslational stage to stabilize TMD1 (Loo *et al*., 2013; Kleizen *et al*., 2021; Fiedorczuk and Chen, 2022a, 2022b), while elexacaftor binds during the second stage, after translation is completed (Fiedorczuk and Chen, 2022b).

Despite these remarkable advances in our understanding of CFTR structure, folding, and correction, our understanding of the mechanism by which CFTR-F508del is triaged and degraded by ERAD remains rudimentary because of the lack of a systematic, genome-wide investigation. Since 2001, at least seven ubiquitin E3 ligases, CHIP/STUB1 (Meacham *et al*., 2001), FBXO2 (Ramachandran *et al*., 2016), NEDD4L (Caohuy *et al*., 2009), RNF185 (Khouri *et al*., 2013), GP78/AMFR (Ballar *et al*., 2010), RMA1/RNF5 (Younger *et al*., 2006; Tomati *et al*., 2015; Sondo *et al*., 2018; Brusa *et al*., 2023), and HRD1 (Ballar *et al*., 2010; Ramachandran *et al*., 2016) have been implicated in CFTR-F508del degradation, yet the relative contribution of each to CFTR-F508del degradation is unknown. Because disruption of CFTR-F508del ERAD can synergize with and/or complement pharmacological correction (Sondo *et al*., 2018; Borgo *et al*., 2022; Brusa *et al*., 2022, 2023), a detailed and comprehensive mechanistic understanding of CFTR ERAD is warranted.

There are two non-mutually exclusive explanations for the lack of a consensus mechanism for CFTR-F508del ERAD: 1) the core machinery has not been identified because the relevant genes were not included in previous candidate-based studies or small-scale siRNA screens, or 2) the ERAD systems that triage CFTR-F508del are so heavily buffered that no single system or set of systems prevails. To distinguish between these two possibilities, we performed a genome-wide single-gene CRISPR/Cas9 knockout screen using a quantitative, fluorescence-based readout of CFTR-F508del stability to comprehensively interrogate CFTR-F508del ERAD. This screen identified RNF5 as the top ERAD E3 ligase hit, with no other single E3 showing a comparable magnitude of effect. Yet, *RNF5* disruption only modestly stabilized CFTR-F508del. Because the genome-wide library contained guides targeting all known E3 ubiquitin ligases, these findings lead us to conclude that F508del ERAD is heavily buffered. To understand the molecular basis of this buffering, we used double-knockouts and sensitized CRISPR knockout screens to determine that the RNF5 ortholog, RNF185, acts redundantly to buffer CFTR-F508del degradation. Importantly, knocking out ERAD components identified in our screens did not increase trafficking to the PM, suggesting that, in the absence of correctors, CFTR-F508del is unable to adopt a conformation that is competent to escape the ER and traffic to the PM, even if its degradation is blocked. Gene-drug interaction analyses with tezacaftor and elexacaftor support a model in which correctors shift the conformational equilibrium toward more native states that are not RNF5/RNF185 substrates.

## RESULTS

### mNG-CFTR K562 cell lines as a model system for interrogating CFTR ERAD

To identify genes that mediate CFTR-F508del ERAD, we adapted a FACS-based CRISPR/Cas9 screening platform developed by the Kopito laboratory to interrogate substrate-specific ERAD (Leto and Kopito, 2019; Leto *et al*., 2019). This approach relies on a doxycycline (dox)-inducible fluorescent reporter encoded by an mRNA lacking a polyadenylation sequence, thereby enabling rapid, selective transcriptional shutoff of the reporter mRNA after dox washout (Leto and Kopito, 2019; Leto *et al*., 2019). Transcriptional shutoff improves the dynamic range of protein degradation screens because the difference in fluorescence between cells carrying sgRNAs that stabilize the reporter and those with sgRNAs that have no effect is greater after reporter expression is shut off and the majority of the reporter has been degraded (Leto *et al*., 2019). While translational shutoff with cycloheximide or emetine allows for more precise temporal control of translation, dox washout enables screens to be performed without the pleiotropic impact of blocking all protein synthesis. Furthermore, the dox-inducible system enables propagation of cells in the absence of transgene expression, thereby avoiding adaptive changes arising from chronic expression of the CFTR-F508del reporter.

To establish reporter cell lines for the CRISPR knockout platform, dox-inducible mNeonGreen (mNG)-CFTR-F508del and mNG-CFTR-WT transgenes were integrated into a K562 cell line expressing the reverse tetracycline-controlled transactivator, rtTA, which binds to and activates Tet-ON promoters in the presence of dox (Das *et al*., 2016). Clonal reporter cell lines expressing mNG-F508del or mNG-WT that exhibited normally distributed fluorescence (**Figure 1A**) with similar median fluorescence intensities (MFI) and degradation kinetics to those of the parental populations were isolated. mNG-F508del cells had a steady-state MFI that was ∼3.5 fold lower than mNG-WT clonal cells (**Figure 1B**), consistent with published data demonstrating that ERAD clearance of CFTR-F508del decreases steady-state CFTR expression (Ward *et al*., 1995; Kälin *et al*., 1999; Chung *et al*., 2016). mNG-F508del was only observed in the immature, core-glycosylated, ER “Band B” glycoform while mNG-WT was observed both in the ER glycoform and in the mature, complex “Band C” glycoform (**Figure 1C**). mNG-F508del and mNG-WT degradation kinetics were assayed by flow cytometry after translational shutoff with the elongation inhibitor, emetine (**Figure 1D**). Degradation kinetics were modeled using a one-phase decay equation (Y = [Y_0_ – Plateau]e^-kx^ – Plateau), which assumes the substrate exists in two states - a short-lived species that follows first-order decay kinetics (Y = Y_0_e^-kx^) and a long-lived species that does not degrade (an asymptotic “plateau”). Although this model oversimplifies the complexity of CFTR turnover, the model is an excellent fit for the CFTR kinetics data we collected in this study, regardless of the chemical or genetic perturbation (median r^2^ of kinetics data in **Figures 1-4** = 0.997). Using the one-phase decay model, we found that the majority of mNG-F508del was degraded with a half-life of 34 minutes (**Figure 1D, Figure S1A**), which is in line with previous kinetics studies of transiently expressed untagged F508del (Ward and Kopito, 1994; Gelman *et al*., 2002). By contrast, ∼40% of mNG-WT was degraded with a half-life of 44 minutes while ∼60% achieved a stable, asymptotic species, consistent with the previous reports showing that a fraction of newly synthesized CFTR-WT is degraded by ERAD before it attains a native conformation that matures to the PM (Ward and Kopito, 1994; Ward *et al*., 1995). Thus, our mNG-tagged CFTR K562 reporter cells recapitulate the classic degradation and maturation phenotypes for CFTR-F508del and CFTR-WT.

**Figure 1:**
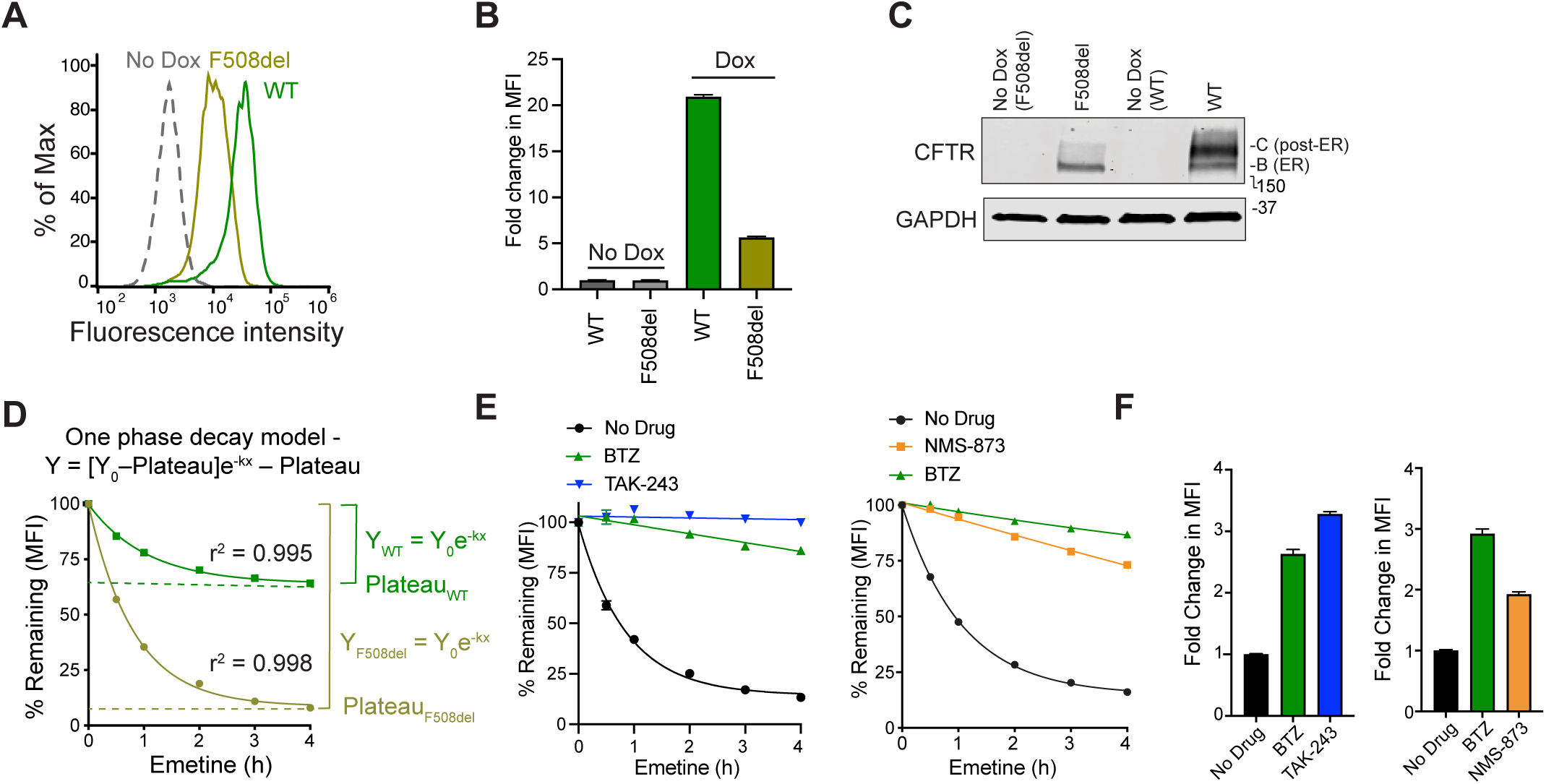
mNG-CFTR K562 cell lines as a model system for interrogating CFTR ERAD. (**A-B**) mNG-CFTR-F508del reporter cells have a lower median fluorescence intensity (MFI) than mNG-CFTR-WT reporter cells. (**A**) Example steady-state fluorescence distributions of clonal mNG-F508del and mNG-WT reporter cell lines after administration of 1 μg/ml doxycycline [dox] for 16 hours and (**B**) quantification of normalized MFI for mNG-F508del and mNG-WT cell lines (n = 3 biological replicates, error bars = SD). (**C**) mNG-F508del and mNG-WT recapitulate CFTR biochemical phenotypes. Immunoblots of reporter cells with and without doxycycline induction depicting complex-glycosylated, post-ER “Band C” and immature, core-glycosylated ER-associated “Band B” glycoforms. MW markers are indicated. (**D**) One phase decay modeling of mNG-F508del degradation. A one phase decay equation was used to model mNG-F508del and mNG-WT degradation kinetics after translational shutoff with translation inhibitor emetine. Percent remaining calculated from MFI using flow cytometry (n = 3 biological replicates, error bars not shown due to small SD between replicates, curve = one-phase decay). See **Figure S1A** for quantification of half-life and plateau data. (**E**) Effect of TAK-243, BTZ, and NMS-873 on mNG-F508del degradation kinetics. As in Figure 1D, kinetics of mNG fluorescence decay following translation shutoff. Fold change in MFI after 3 hours of proteasome inhibition with 1 μM bortezomib (BTZ) (left and right), E1 inhibition with 2 μM TAK-243 (left), and VCP inhibition with 20 μM NMS-873 (right). (**F**) Effect of TAK-243, BTZ, and NMS-873 on steady-state levels of mNG-F508del. As in Figure 1E, measurements taken at t = 0 prior to the addition of emetine.

To confirm that mNG-F508del and mNG-WT are targets of the UPS, degradation kinetics experiments were performed in the presence of pharmacological inhibitors of the proteasome (Bortezomib, BTZ), the E1 ubiquitin-activating enzyme (TAK-243), and the p97/VCP AAA+-ATPase (NMS-873). All three inhibitors stabilized mNG-F508del (**Figure 1E**) and mNG-WT (**Figure S1B**). The fold-changes in protein expression were greater for mNG-F508del (**Figure 1F**) than mNG-WT (**Figure S1C**), underscoring that the F508del mutation diverts a greater fraction of CFTR to the UPS. The E1 inhibitor, TAK-243, fully stabilized mNG-WT and mNG-F508del degradation (**Figure 1E, Figure S1B**), suggesting that CFTR degradation is entirely ubiquitin dependent. TAK-243 led to a greater increase on CFTR steady-state and degradation kinetics than the proteasome inhibitor, bortezomib (BTZ), suggesting that some CFTR molecules are degraded through a ubiquitin-dependent, proteasome-independent process. We speculate that a small fraction of CFTR-F508del escapes ERAD and is subsequently degraded through a ubiquitin-dependent endolysosomal process known to target temperature-rescued CFTR-F508del at the PM (Okiyoneda *et al*., 2010, 2018; Taniguchi *et al*., 2022). VCP inhibition by NMS-873 had a more modest impact on expression levels and degradation kinetics than BTZ or TAK-243, suggesting that not all CFTR molecules undergo VCP-mediated dislocation from the ER membrane. The lack of full stabilization by BTZ or NMS-873 was not due to incomplete inhibition because increasing the concentration of inhibitors did not improve the stabilization of CFTR-F508del (**Figure S1D**) and because other model ERAD substrates using our K562 platform were equally stabilized with NMS-873 and the proteasome inhibitor, MG-132 (Leto *et al*., 2019). Therefore, CFTR-F508del molecules have at least one of three fates: 1) VCP-dependent, proteasome-dependent degradation, 2) VCP-independent, proteasome-dependent degradation, or 3) proteasome-independent, ubiquitin-mediated degradation.

In sum, validation experiments demonstrate that the K562 mNG-CFTR-F508del reporter recapitulates the quintessential CFTR-F508del phenotypes, providing us with a facile and quantitative model system for interrogating CFTR-F508del ERAD in large-scale CRISPR knockout screens.

### Genome-wide CRISPR analysis reveals that CFTR-F508del degradation is highly buffered

We conducted genome-wide screens of CFTR-F508del degradation using a 10-guide per gene human CRISPR/Cas9 knockout sgRNA library (Morgens *et al*., 2017) and used casTLE analysis (Morgens *et al*., 2016) to compare the sgRNA frequencies in the sorted top 5% of the mNG fluorescence distribution with those in the unsorted population (**Figure 2A**). Of the 20,528 genes assayed in our screen, 207 genes were high-confidence significant hits (FDR < 1%, **Figure 2B**, **Table S1, Figures S2A-B**). As previously reported (Leto *et al*., 2019), mRNA decay genes like *XRN1* and *XRN2* were among the top hits because our transcriptional shutoff assay requires the activity of exonucleases to degrade the reporter transcript (**Figure S2C**).

**Figure 2:**
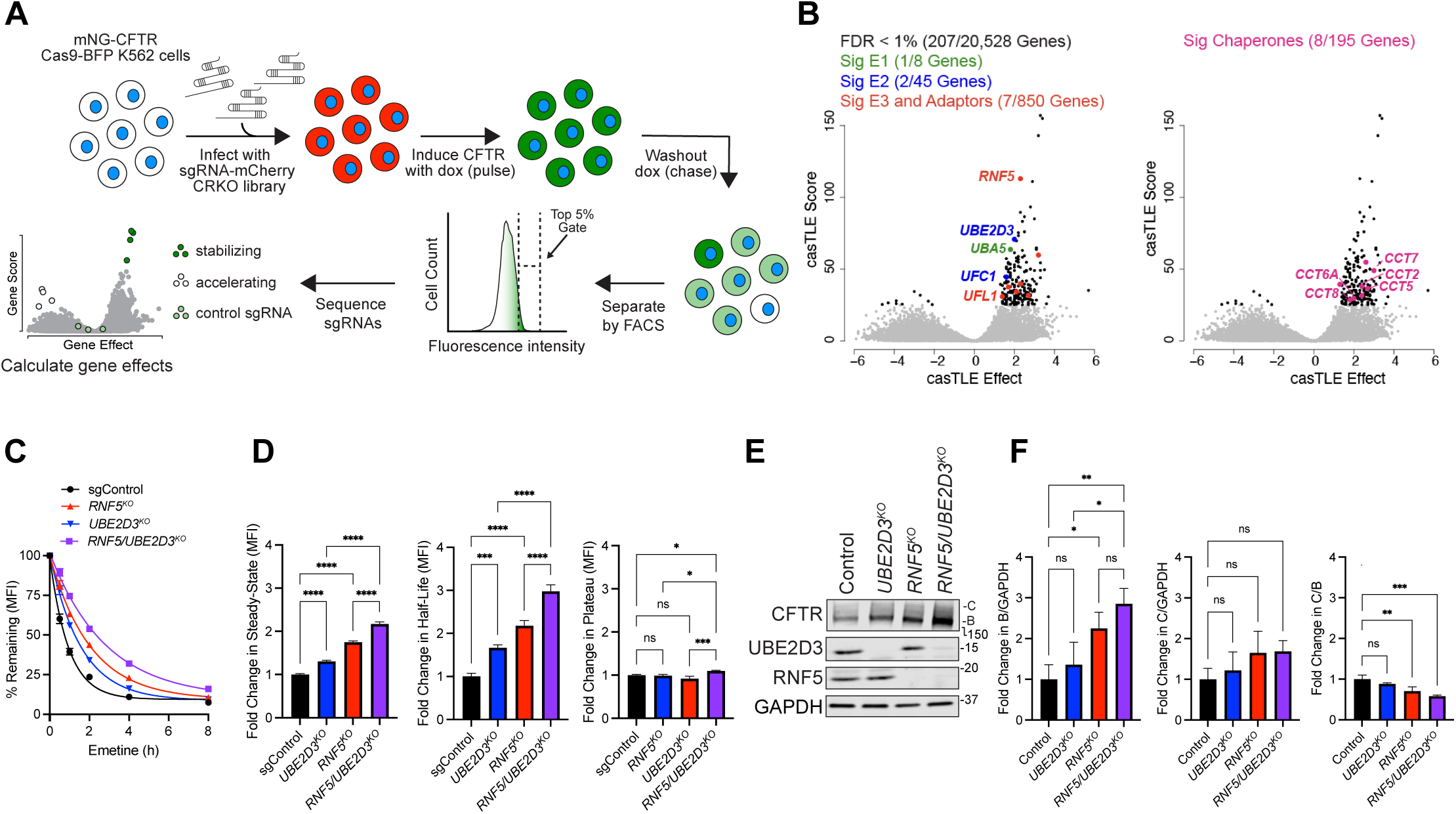
Genome-wide CRISPR analysis reveals that CFTR-F508del degradation is highly buffered. (**A**) Schematic of genome-wide screening method. (**B**) Volcano plots of single knockout CRISPR screen. casTLE analysis of genome-wide screens with mNG-F508del (n = 2 biological replicates). Grey = genes observed in screens. Black = significant genes (FDR < 1%). Red = significant E3 ubiquitin ligases and adaptors. Blue = significant E2 ubiquitin conjugating enzymes. Green = significant E1 ubiquitin activating enzymes. Pink = significant chaperones and co-chaperones. See **Figure S2A** for doxycycline washout curves for screens, **Figure S2B** for correlation between replicates, and **Table S1** for complete casTLE analysis. (**C-F**) Validation of RNF5 and UBE2D3 hits. Pooled knockout mNG-F508del cell lines were generated using Cas9 RNP nucleofection with three sgRNAs per gene. sgControl is a control guide targeting the *AAVS1* safe-harbor locus (* = p < 0.05, ** = p < 0.01, *** = p < 0.001, **** = p < 0.0001, one-way ANOVA with Tukey’s multiple comparisons test). (**C**) Effect of gene disruption on mNG-F508del degradation kinetics, as in Figure 1D. (**D**) Quantification of data from (**C**). (Left) fold change in steady-state MFI at t = 0 prior to the addition of emetine. (Center) half-life and (right) plateau as in Figure 1D & **Figure S1A**. (**E**) Immunoblot of mNG-F508del expression and knockout efficiency as in Figure 1C. (**F**) Quantification of data from Figure 2E (n = 3 biological replicates imaged on the same immunoblot).

Given that CFTR-F508del degradation is entirely dependent on ubiquitin conjugation (**Figure 1E**), we focused our analysis on identifying the key components of the ubiquitin modification machinery that contribute to CFTR-F508del degradation. One E1 ubiquitin/ubiquitin-like activating enzyme (*UBA5*), 2 E2 ubiquitin/ubiquitin-like conjugating enzymes (*UBE2D3*, *UFC1*), and 7 E3 ubiquitin ligases/ligase adaptors passed our significance filter (FDR < 1%, **Figure 2B, Table S1**). RNF5, a membrane-bound E3 ligase previously implicated in CFTR-F508del degradation (Younger *et al*., 2006; Khouri *et al*., 2013; Tomati *et al*., 2015), was the top E3 ligase, yet none of the other six E3 ligases/ligase adaptors previously implicated in CFTR-F508del degradation was significant (**Figure 2B, Table S1**). Most of the remaining E3 ligases and adaptors identified (*PPIL2, KCTD5, PHF5A, RTEL1, CDC16)* are annotated to be involved in transcription or cell growth with no clear relationship to CFTR-F508del degradation or ERAD.

The top E2 hit was UBE2D3, a ubiquitin conjugating enzyme previously implicated in CFTR-F508del degradation at the PM (Okiyoneda *et al*., 2010) but also known to interact with RNF5 (Tsai *et al*., 2022). A second E2, UFC1, is the conjugating enzyme for the ubiquitin-like protein, UFM1. While several other components of the UFMylation pathway were also identified in our screen (*UBA5*, *UFC1*, *UFL1*), we found that targeted disruption of these genes did not influence CFTR-F508del turnover but, instead, increased CFTR-F508del transcript abundance (**Figure S2D-E**).

Strikingly, 5 out of 10 genes encoding the tailless complex polypeptide ring complex (TriC) subunits were significantly enriched (**Figure 2B**), suggesting a role of this cytoplasmic chaperonin in triaging and directing CFTR-F508del to the proteasome for degradation. TriC subunits were previously reported to be part of the core interactome for CFTR-F508del and CFTR-WT, and association of these components with CFTR-F508del was reduced when folding of cytosolic NBD1 was promoted via shifting cells to low temperatures or dosing cells with a pharmacological inhibitor of histone deacetylases (Pankow *et al*., 2015). TriC may be involved in triaging the unfolded cytosolic domains of CFTR-F508del. The other 3 chaperones/co-chaperones identified (*RUVBL1*, *RUVBL2*, and *TSC1,* **Table S1**) are annotated to be involved in chromatin remodeling (Puri *et al*., 2007) and cell growth (Tee *et al*., 2002), respectively and are unlikely to contribute directly to CFTR-F508del folding or triage.

The casTLE gene effect, a measure of sgRNA enrichment in the sorted population, and the casTLE gene score, a measure of the confidence of the casTLE gene effect, were considerably lower for hits observed in the mNG-F508del screen than for hits in previous screens with model ERAD substrates that are degraded by linear ERAD pathways (**Figure S2F**), suggesting that multiple parallel or redundant pathways contribute to CFTR-F508del ERAD. Consistent with this interpretation, knocking out the genes encoding the top E3 (RNF5) and E2 (UBE2D3) had only modest impacts on steady-state mNG-F508del levels and degradation kinetics (**Figure 2C**); disrupting both genes simultaneously exhibited additive effects on degradation kinetics and steady-state levels for both mNG-F508del and mNG-WT (**Figure 2C-D, Figure S2G**). Disrupting *RNF5* and *UBE2D3* singly or in combination increased the levels of Band B but not Band C of the mNG-F508del reporter (**Figure 2E-F**), suggesting that ERAD targets a CFTR-F508del conformer that is not competent to exit the ER. Our findings suggest that RNF5-mediated ubiquitylation is not uniquely mediated by UBE2D3 and that other E3s and E2s contribute redundantly with UBE2D3 and RNF5 (**Figure 2C-D, S2G**). The comparatively low casTLE gene effect scores and correlations between screen replicates observed for our top hits reflect the modest impact to mNG-CFTR-F508 degradation achievable by knocking out any single gene in the human genome. Taken together with the finding that exposure to the ubiquitin E1 inhibitor, TAK-243, and the proteasome inhibitor, BTZ results in near-complete CFTR-F508del stabilization, our results lead to the conclusion that CFTR-F508del degradation is mediated by genetically redundant, ubiquitin-dependent, proteasome-dominated processes.

### Sensitized CRISPR screens identify genetically redundant ubiquitin conjugation machinery for CFTR-F508del ERAD

To identify redundant UPS modules that mediate CFTR-F508del degradation, we conducted parallel CRISPR knockout screens in *RNF5^KO^*, *UBE2D3^KO^*, and *RNF5/UBE2D3^KO^* reporter cell lines in replicate with an sgRNA sublibrary composed of guides targeting ∼2,000 genes involved in the <u>ub</u>iquitin, <u>a</u>utophagy, and lysosomal (UBAL) degradation pathways (**Table S2**). This custom-designed sublibrary targets all known ubiquitin conjugation and deconjugation enzymes, ubiquitin-like conjugation systems, and proteasome components, thereby enabling rapid and comprehensive functional genomics interrogation of the ubiquitinome. As with the genome-wide screens, RNF5 and UBE2D3 were the top E3 and E2 hits, respectively, in the control single-gene UBAL sublibrary screens, validating our sublibrary screening approach. Other than RNF5, no other E3 ligases previously implicated in CFTR-F508del ERAD were identified as significant hits (FDR < 1%), and at least 11 of the other 13 E3 ligases identified in the control screen appear to have roles in transcription, DNA damage signaling, and cell growth.

Different profiles were observed when the UBAL library was screened in “sensitized” genetic backgrounds harboring deletions of *RNF5*, *UBE2D3,* or both genes together (**Figure 3A**, **Table S3**). When the sensitized screen was conducted in cells that lack *RNF5* (i.e., *RNF5^KO^* or *RNF5/UBE2D3^KO^*), its close ortholog, RNF185, emerged as the top E3 ligase. RNF185 shares 70% amino acid identity with RNF5 and has previously been implicated in turnover of ER membrane proteins including CFTR-F508del (Khouri *et al*., 2013; Weijer *et al*., 2020). Knocking out *RNF185* alone negligibly increased reporter half-life, indicting that, when RNF5 is present, it does not contribute significantly to CFTR-F508 turnover and corroborating the absence of *RNF185* from the single-gene genome-wide and control sgControl UBAL screens (**Figure 3B**, **Table S3**). By contrast, knocking out both *RNF185* and *RNF5* together synergistically increased reporter half-life (**Figure 3B**) and Band B levels (**Figure 3C**), suggesting that RNF185 acts redundantly with RNF5 to degrade CFTR-F508del. In addition to RNF185, a second E3, GP78/AMFR and its associated E2, UBE2G2 (Chen *et al*., 2006), were found to be weak, but significant hits in the *RNF5^KO^* screen (**Figure 3A**), suggesting that this E3 could contribute to CFTR-F508del degradation.

**Figure 3:**
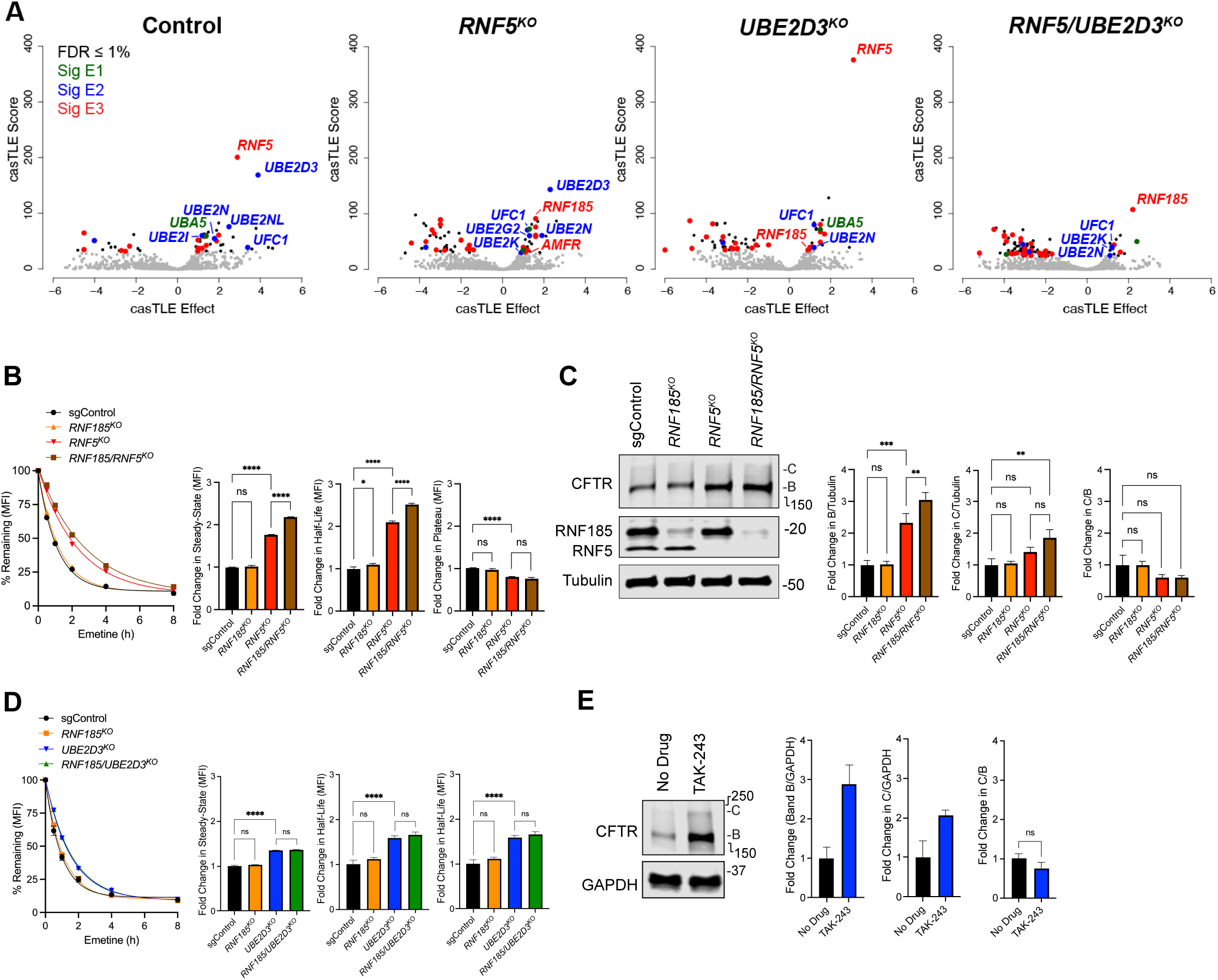
Sensitized CRISPR screens identify genetically redundant ubiquitin conjugation machinery for CFTR-F508del ERAD. (**A**) casTLE analysis of sensitized screens with the UBAL sublibrary. Grey = genes observed in screen. Black = significant genes (FDR < 1%). Red = significant E3 ubiquitin ligases. Blue = significant E2 ubiquitin conjugating enzymes. Green = significant E1 ubiquitin activating enzymes. n = 2 biological replicates per genotype. (**B-D**) Knocking out *RNF185* and *RNF5* simultaneously has a synergistic effect on mNG-F508del half-life (**B**) but does not increase maturation (**C**). (**D**) mNG-F508del has an equivalent half-life in *RNF185/UBE2D3^KO^* and *UBE2D3^KO^* backgrounds. Methods and analyses are the same as in Figure 2C-F. (**E**) Inhibiting ubiquitylation does not increase mNG-F508del maturation. Immunoblots of mNG-CFTR-F508del cells treated with 2 μM TAK-243 for 3 hours after dox induction. Methods and analyses are the same as in Figure 2E-F.

The sensitized screens also provided insights into the identities of and possible interactions among E2 ubiquitin conjugating enzymes that contribute to RNF5-and RNF185-mediated CFTR-F508del ERAD. Notably, the sensitized screen in *RNF5^KO^* cells identified both UBE2D3 and UBE2K (**Figure 3A**), E2s which were both previously implicated in RNF185-dependent degradation of the model single-pass transmembrane ERAD substrate CPY51A1 (Weijer *et al*., 2020). Although RNF185 was a modest hit in the pooled *UBE2D3^KO^* cells (**Figure 3A**), mNG-F508del exhibited indistinguishable degradation kinetics in *RNF185*/*UBE2D3^KO^* cells as in *UBE2D3^KO^* cells (**Figure 3D**), suggesting that the genes could act in the same pathway. Given that UBE2D3 was the only E2 identified in our genome-wide screens yet knocking out *UBE2D3* and *RNF5* was additive (**Figure 2C-F**), we propose that UBE2D3 can function as an E2 for both RNF5 and RNF185 in the context of CFTR-F508del degradation. While our CRISPR knockout data from *RNF5/UBE2D3^KO^* cells suggested that RNF185-mediated ERAD is not solely dependent on UBE2D3, our attempts to determine if RNF5, RNF185, and UBE2D3 acted together were confounded by the fact that *RNF185/RNF5/UBE2D3^KO^* triple knockouts exhibited a severe growth defect, and we were unable to culture stable pools or isolate clones. Although we speculate that these three proteins overlap to mediate an essential ERAD process, we cannot rule out the possibility that knocking out the three genes is lethal through some unrelated mechanism.

We also identified two E2s in our control UBAL screens that are involved in ubiquitin-like conjugation pathways, *UBE2I* and *UFC1*, which encodes E2s in the SUMOylation, and UFMylation pathways, respectively (**Figure 3A**). Although other SUMOylation machinery was not identified in our screens, it is noteworthy that SUMO modification has been previously linked to CFTR modulation (Ahner *et al*., 2016). Finally, UBE2N was a significant hit in all the UBAL sublibrary screens (**Figure 3A**). This E2, which selectively conjugates Lys-63 linked ubiquitin to ubiquitin “primed” substrates, is implicated in ubiquitin chain diversification in a broad range of contexts including cell cycle regulation and DNA damage response (Stewart *et al*., 2016). One report links UBE2N-dependent K63 chains to ERAD (Wolf *et al*., 2021) while another reported that UBE2N interacts with RNF5 to ubiquitylate the JNK-associated membrane protein (JKAMP) in a process that out-competes RNF5-mediated ERAD of CFTR-F508del (Tcherpakov *et al*., 2009).

Collectively, our sensitized screens demonstrate that CFTR-F508del ERAD is highly buffered, with RNF185 acting redundantly with RNF5. Moreover, no single E2 was shown to exhibit a synergistic genetic interaction with UBE2D3 in our *UBE2D3^KO^* screen, suggesting that at least three redundant E2s are involved (**Figure 3A**, **Table S3**). Importantly, none of the single or double knockouts of known ERAD machinery increased CFTR-F508del trafficking to the PM (**Figure 2E-F**, **Figure 3C**), and indeed, dosing cells with the general E1 inhibitor, TAK-243, does not increase the Band C/B ratio (**Figure 3E**), (Borgo *et al*., 2022). Thus, in the absence of correctors, CFTR-F508del is unable to attain a folding state that is competent to escape the ER or to traffic to the PM, even in the near-absence of degradation by ERAD.

### Correctors promote CFTR-F508del maturation into ERAD-resistant folding states

Structural analysis of CFTR-F508del complexes with tezacaftor and elexacaftor alone or in combination suggest that these correctors promote folding by stabilizing more native-like sequential folding states(Fiedorczuk and Chen, 2022b). We propose a model in which newly synthesized CFTR molecules transit through three discrete sequential folding states in the ER membrane (B_1_, B_2_, and B_3_) that are in equilibrium with one another (**Figure 4A**) with the earliest of these (B_1_) targeted for degradation via RNF5/RNF185. In the absence of correctors, the net flux for CFTR-F508del is shifted towards B_1_ (denoted by thicker arrows in **Figure 4A**), ensuring that most mutant molecules fail to mature and are instead degraded. Our model posits that correctors promote folding by binding to and stabilizing later folding states, B_2_ and B_3_. Because corrector binding is reversible, some ERAD occurs even in their continued presence, until the folded molecule escapes from the ER.

We found that degradation (**Figure 4B**) and maturation (**Figure 4C**) of CFTR-F508del was negligibly increased in the presence of tezacaftor (T) and elexacaftor (E) alone but was substantially increased in their combined presence, confirming that these drugs act synergistically on our reporters in K562 cells. While tezacaftor modestly increased the half-life of CFTR-F508del, elexacaftor did not significantly alter half-life but increased the long-lived “plateau” species (**Figure 4B**). Taken together, these data support a model in which tezacaftor stabilizes an early folding state (B_2_) while elexacaftor stabilizes a later state (B_3_) that can exit the ER and traffic to the PM. Indeed, the NBD1 domain, which contains the F508del mutation and the ER exit code (Wang *et al*., 2004) is unstructured in lumacaftor-bound CFTR-F508del while it is structured in the elexacaftor-and ET-bound states (Fiedorczuk and Chen, 2022b). In the presence of elexacaftor alone, the B_1_ state is not stabilized and ERAD is not inhibited, but the fraction of CFTR-F508del molecules that achieve the B_2_ conformation are stabilized by binding to the drug and can exit the ER. However, the addition of tezacaftor increases the amount of B_2_ available for elexacaftor to bind, and the drugs synergize to increase CFTR-F508del trafficking and function.

**Figure 4:**
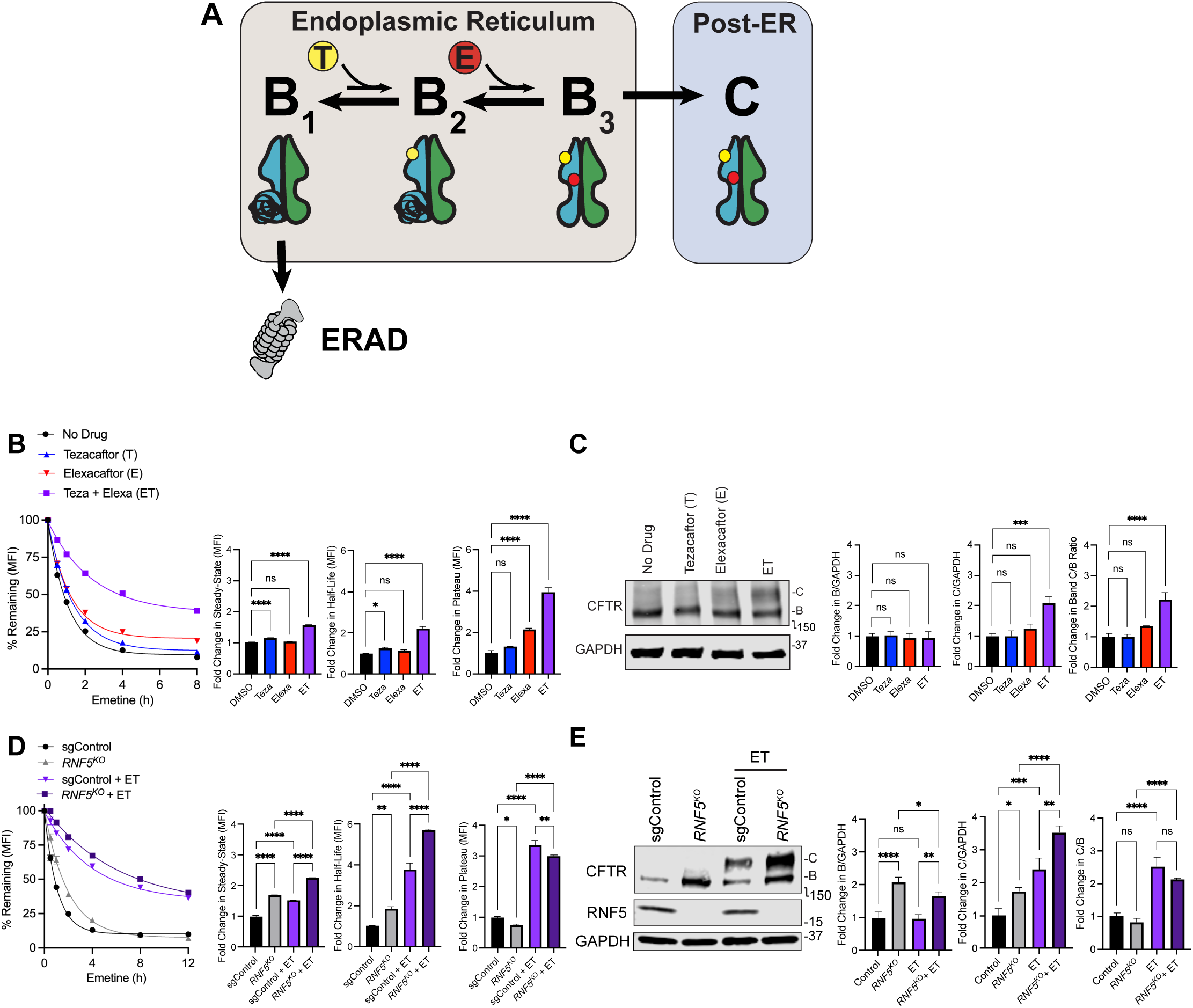
Correctors promote CFTR-F508del maturation into ERAD-resistant folding states. (**A**) Model of CFTR-F508del folding, correction, and triage. Sequential CFTR-F508del folding states are in equilibrium at the ER and converting between states is increased by corrector binding. The effect of the F508del mutation on the equilibrium between core-glycosylated states B_1_-B_3_ is indicated by the thickness of forward and reverse arrows. B_1_: Nascent CFTR-F508del with unstructured NBD1. This folding state is targeted by ERAD and is incapable of exiting the ER. B_2_: more native-like folding state that is not an ERAD substrate. Binding of tezacaftor to B_2_ lowers probability of converting back to B_1_. B_3_: later folding state that is not an ERAD substrate and has the capacity to bind to elexacaftor, which slows the rate of conversion back to B_2_. B_3_ is near-native with a structured NBD1 and can exit the ER. C: complex-glycosylated folding-state in post-ER compartments. While binding of correctors changes the equilibrium constants between states, knocking out ERAD components does not. Structural cartoons are adapted from (Fiedorczuk and Chen, 2022b). (**B-E**) Inhibiting ERAD increases the amount of mNG-F508del available for correction. Cells were administered tezacaftor (VX-661, 5 μM) and/or elexacaftor (VX-445, 5 μM) for 16 hours. Elexacaftor and tezacaftor synergistically increase mNG-F508del half-life (**B**) and maturation (**C**). Knocking out *RNF5* increased the effect of ET on mNG-F508del half-life (**D**) and maturation (**E**). Methods and quantification are as in Figure 2C-F.

As previously reported (Capurro *et al*., 2021), tezacaftor modestly increases the half-life of CFTR-F508del (**Figure 4B**). This stabilization could be due to tezacaftor interfering with ERAD or increasing the probability of CFTR-F508del occupying an ERAD-resistant folding state. To distinguish between these possibilities, we performed epistasis experiments with tezacaftor and *RNF5* knockout. If tezacaftor interferes with ERAD, we would expect the half-life of mNG-F508del would be the same in *RNF5^KO^* cells with or without tezacaftor. However, we observed that the effect of combining *RNF5* disruption with tezacaftor on mNG-F508del half-life and maturation was additive (**Figure S3A-B**). Thus, tezacaftor does not prevent RNF5-mediated ERAD but rather increases the likelihood that CFTR-F508del achieves a folding state that is RNF5-resistant. Given that the phenotypes are additive, we conclude that tezacaftor increases the probability that CFTR-F508del transitions from the B_1_ to the B_2_ folding state. However, because B_2_ and B_1_ states are in equilibrium, CFTR-F508del is still rapidly degraded in the presence of tezacaftor when B_2_ reverts to B_1_.

Our data with tezacaftor and *RNF5^KO^* cells suggest that knocking out ERAD components does not increase the probability that CFTR-F508del can convert to the later folding ensembles, but rather increases the total amount of CFTR-F508del in the B_1_ state available for correction. To test this model, we performed gene-drug epistasis experiments with *RNF5^KO^* cells and elexacaftor. If inhibiting ERAD increases the probability that CFTR-F508del transitioned to a later folding state, knocking out *RNF5* should phenocopy tezacaftor and the gene-drug interaction would be synergistic. However, we observed that knocking out *RNF5* had additive effects with elexacaftor on slowing CFTR-F508del degradation kinetics (**Figure S3C**) and increasing maturation (**Figure S3D**). Thus, knocking out *RNF5* increases the amount of B_1_ available for correction but does not increase the probability of conversion to B_2_ thereby making more CFTR-F508del available for elexacaftor to convert to B_3_. These data lead us to conclude that inhibiting ERAD increases the amount of unfolded, trafficking incompetent B_1_ intermediates that are available for correction. Indeed, knocking out *RNF5* increased the effect of ET on mNG-F508del half-life (**Figure 4D**) and maturation (**Figure 4E**), by 52% and 46%, respectively. Likewise, knocking out *UBE2D3* was additive with the combination of tezacaftor and elexacaftor on degradation kinetics (**Figure S3E-G**) and maturation (**Figure S3H**).

## DISCUSSION

### CFTR-F508del ERAD is mediated through multiple quality control pathways

In this study, we employed genome-wide CRISPR loss-of-function screens to investigate the mechanism of CFTR-F508del ERAD. We found that no single E3 ligase was responsible for CFTR-F508del ERAD as is the case with many other well-defined ERAD substrates (Menzies *et al*., 2018; Leto *et al*., 2019). The small gene effects observed in our full-scale CRISPR/Cas9 knockout screens underscore the historic challenge of defining CFTR-F508del ERAD using candidate-based studies and siRNA screens and support the conclusion that the PQC network that mediates CFTR-F508del ERAD is highly buffered, involving redundant and overlapping pathways.

The sensitized screens with *RNF5^KO^* and *RNF5/UBE2D3^KO^* cells reveal that *RNF185* can compensate for the loss of *RNF5*, but the observation that disruption of *RNF185* in otherwise wild-type cells does not affect CFTR-F508del turnover supports the view that these two orthologous E3 ligases back each other up. Uncovering the network mediating CFTR-F508del ERAD using sensitized or dual-knockout screens may prove difficult due to the possibility that disruption of redundant core PQC nodes can be synthetically lethal. Consequently, CRISPR/Cas9 screening technologies, such as CRISPRi (Gilbert *et al*., 2014), that transiently knock down genes may be a more productive route for fully describing the highly buffered cellular pathways that triage CFTR-F508del.

A comprehensive understanding of CFTR-F508 triage is further complicated by the fact that ERAD, like other UPS mediated quality control processes, is best viewed as an integrated network or system rather than a linear pathway. Combinations of E3s, acting together with their cognate E2s, work sequentially or in tandem to build topologically complex ubiquitin chains that specify temporally and spatially complex processes in addition to proteasomal degradation (Kolla *et al*., 2022; Ohtake, 2022). Previous CRISPR/Cas9 screens have been instrumental in identifying E2/E3 combinations that build branched chains with complex linkage topologies on diverse ERAD clients (Liu *et al*., 2017; Leto *et al*., 2019). Our identification of GP78/AMFR and its cognate E2, UBE2G2, in the *RNF5^KO^* sensitized screen may be an example, as this E3 was previously reported to function as an E4 that acts downstream of RNF5 in CFTR-F508del ERAD (Morito *et al*., 2008), possibly to increase efficiency of extraction by p97/VCP (Vij *et al*., 2006; Ballar *et al*., 2010).

Knocking out *UBE2D3* had an additive effect with disrupting *RNF5* on CFTR-F508del degradation, although we did not uncover a second E2 with a comparable gene effect score in our single knockout screens with *UBE2D3^KO^*. Therefore, we propose that UBE2D3 can partner with both RNF5 and RNF185, and that other E2s can compensate for its function when it is knocked out. This idea is supported by a previous study reporting that the role of RNF5 in degrading a Lamin B receptor disease variant is partly dependent on UBE2D3 (Tsai *et al*., 2022). UBE2D3 and UBE2K were shown to be partially redundant in RNF185-mediated degradation of the membrane domain of the lanosterol demethylase, CPY51A1 (van de Weijer *et al*., 2020). Given our identification of UBE2K in the *RNF5/UBE2D3^KO^* and *RNF5^KO^*, but not the *UBE2D3^KO^* screens (**Table S3**), supports the argument that UBE2K acts redundantly with UBE2D3 to mediate RNF185 triage of CFTR-F508del. Further candidate-based studies or genome-wide sensitized screens will be necessary to determine if other members of the RNF185 complex required for CPY51A1 ERAD, like membralin and TMUB1/2 (van de Weijer *et al*., 2020) are also required for CFTR-F508del ERAD.

The effects on CFTR-F508del half-life and state-state levels that we observe following CRISPR/Cas9 disruption of *RNF5* and *RNF185* alone and in combination were more modest than those previously reported when using RNA silencing in HEK293 cells transiently expressing CFTR-F508del (Khouri *et al*., 2013). Given that K562 cells contain 4 and 2 copies of *RNF5* and *RNF185*, respectively (Zhou *et al*., 2019), it is likely that our knockout population was a heterogeneous pool that contained some cells harboring one or more unedited loci. Therefore, we cannot rule-out that the more modest phenotype was due to residual expression.

### Tezacaftor and elexacaftor promote sequential CFTR-F508del folding states at the ER

Our functional genomics platform permitted rapid, quantitative gene-drug interaction experiments that define the relationship between ERAD and corrector drugs, confirming structural models (Fiedorczuk and Chen, 2022b) and biochemical insights (Tomati *et al*., 2015; Sondo *et al*., 2018; Brusa *et al*., 2023). Our data support a model in which the equilibrium between CFTR-F508del folding intermediates in the ER is shifted away from ERAD susceptibility by binding to correctors. Because inhibiting ERAD does not promote trafficking but increases the amount of CFTR available for correction, proteostasis modulators that inhibit ERAD components like RNF5 will prove to be most useful in combination with pharmacological chaperones that stabilize more native states. Combinatorial approaches with small molecules targeting different ERAD components may prove more effective than single treatments, and indeed, the homology between RNF5 and RNF185 could be exploited to design small molecules that inhibit both proteins simultaneously.

## MATERIALS AND METHODS

### Cloning of CFTR reporter plasmids

mNeonGreen-CFTR-WT and mNeonGreen-CFTR-F508del were cloned into pMCB497-pTRE-polyA(-)-pPGK-blast^R^ vector backbone (Leto *et al*., 2019) using the NEB HiFi Assembly mix (NEB, Ipswitch, MA). CFTR was amplified from pShu-eGFP-CFTR-WT and pShu-eGFP-CFTR-F508del (gift from Jon Hanrahan, McGill University) while mammalian codon-optimized mNeonGreen (mNG) was amplified from a double-stranded DNA gene block (gift from Puglisi Laboratory, Stanford University).

pShu-eGFP-CFTR-WT and pShu-eGFP-CFTR-F508del were sequenced at Sequetech (Mountain View, CA) using an Applied Biosystems 3730/3730*xl*DNA Analyzer (Thermo Fisher Scientific, Waltham, MA) to identify aberrant non-synonymous mutations in the CFTR coding sequences. V470M was included in our F508del construct, as V470M is linked to F508del in CF populations (Vecchio-Pagán *et al*., 2016). Primers carrying the sequence to be corrected were designed, and the plasmid was amplified using Q5 DNA polymerase (NEB). The template plasmid was digested with DpnI (NEB), and the PCR amplicon was purified using the DNA Clean & Concentrator-5 (Zymo Research, Irving, CA). The amplicon was incubated in 10X T4 ligase buffer (NEB) and PNK (NEB) at 37 °C for 1 hour prior to the addition of T4 DNA ligase (NEB). The ligation reaction was incubated at room temperature for 1 hour then transformed into DH5ɑ competent cells.

### Cell culture

HEK 293T cells (ATCC, Manassas, VA) were cultured in DMEM 4.5 g/L glucose, L-glutamine, without sodium pyruvate (Corning, Corning, NY) with 10% FBS (VWR, Radnor, PA). K562 cells were cultured in RPMI medium with L-glutamine with or without phenol red (Corning, Corning, NY). All cells were cultured at 37 °C in 5% CO_2_.

### Small scale lentiviral transduction of K562 cells

HEK293T cells were plated at 20-30% confluency in 2 ml DMEM with 10% FBS in 6-well dishes. The following day, the cells were transfected using TransIT-LT1 (Mirus Bio LLC, Madison, WI) with 0.75 μg third-generation lentiviral packaging mix (pMD2.G (Addgene #12259), pRSV-Rev (Addgene #12253), and pMDLg/pRRE, (Addgene #13351)) (Dull *et al*., 1998) and 0.75 μg lentiviral plasmid. After 24 hours, 2 ml fresh DMEM + FBS was added to each well, and 48 hours later, viral media was filtered through a 0.45 μm PVDF membrane filter (Genesee Scientific, San Diego, CA). Viral media was used immediately or frozen at −80 °C.

K562 cells were spun at 800 x *g* for 5 minutes at room temperature, and 500,000 cells were resuspended in 500 μl viral media. The cells were then spun at 1000 x *g* in a M-20 Microplate Swinging Bucket Rotor (Thermo Fisher Scientific) on a Sorvall Legend XTR centrifuge (Thermo Fisher Scientific) for 2 hours in 12-well plates at 33 °C. The viral media was removed, and the cells were placed in fresh growth media. Cells were placed on selective media 3 days after viral transduction.

### Generation of reporter cell lines for CRISPR knockout screens

Polyclonal K562 cells carrying an rtTA transactivator co-expressed with a G418 resistance marker (pEF1ɑ-TetON Advanced-IRES-G418^R^, (Leto *et al*., 2019)) were transduced with lentivirus generated from pMCB-497-pTRE-mNeonGreen-CFTR-WT-polyA(-)-pPGK-blast^R^ and pMCB-497-pTRE-mNeonGreen-CFTR-F508del-polyA(-)-pPGK-blast^R^. Three days after lentiviral transduction, K562 cells were dosed with 400 μg/ml G418 (Thermo Fisher Scientific) and 7.5 μg/ml blasticidin (Thermo Fisher Scientific) until control cells lacking blasticidin resistance were dead. Reporters were induced with 1 μg/ml doxycycline (Sigma-Aldrich, St. Louis, MO), and mNeonGreen positive cells were sorted at the Stanford Shared FACS facility using a BD FACSAria II Cell Sorter (BD Biosciences, San Jose, CA) equipped with a blue 488 nm laser with a 502 nm long pass mirror and 525/50 nm filter.

Sorted populations were dilution cloned into 96-well plates, and clones were characterized using the protocols detailed in (Leto and Kopito, 2019). Over 200 clones per genotype were analyzed for 1) normal distributions upon dox induction, 2) degradation kinetics identical to that of the polyclonal population, 3) stabilization after treatment with NMS-873, MG-132, or VX-809, and 4) correct glycosylation patterns for CFTR-F508del and CFTR-WT. For the genome-wide knockout screens, reporter cells were transduced with pMH0007-UCOE-pEF1ɑ-Cas9-HA-2xNLS-BFP (Addgene #174162, (Hein and Weissman, 2021)), which contains upstream chromatin opening elements (UCOEs) to prevent Cas9-BFP silencing in K562 cells. Reporter cells were sorted twice for BFP expression with a BD Influx Cell Sorter (BD Biosciences) equipped with a 50 mW violet 405 nm laser and 460/50 nm filter at the Stanford Shared FACS facility.

### Flow cytometry steady-state and degradation kinetics experiments

Fluorescence intensities of mNG-CFTR K562 cells were measured using an Attune NxT Acoustic Focusing Cytometer (Thermo Fisher Scientific) equipped with 405 nm, 488 nm, 561 nm, and 638 nm lasers. K562 cells were gated using FSC-A v SSC-A then single cells were gated using FSC-A v FSC-H. The mNG signal was detected using the 488 nm laser with the 530/30 nm filter (BL1 channel) and gated using BL1-A. Approximately 200 μl K562 cells in growth media were collected and placed on ice, and cells were vortexed and run at 100 μl/min for 10,000 events or 100 μl sample volume. For degradation kinetics experiments, cells were induced with 1 μg/ml doxycycline for 16 hours before 20 μM emetine was added to shut off translation. Median fluorescence intensities (MFI) for singlet K562 cells were computed using the Attune Nxt Software v3.1.2 or FlowJo v10. No dox controls were run in replicate (typically n = 3) as controls for background fluorescence, and the average MFI was subtracted from the MFI at t = n. % remaining was calculated as MFI at t = n minus the MFI of cells without doxycycline divided by the MFI at t = 0 minus the MFI of cells without doxycycline. One-phase decay curves were drawn using GraphPad Prism v9 using the default settings. One curve was drawn per experimental replicate, and the half-life and plateau were calculated, averaged, and normalized to the control.

Flow cytometry degradation kinetics experiments with single and double knockout reporter cells with Cas9-BFP and mCherry-sgRNA transgenes were performed by measuring the median fluorescence intensity of mNG-CFTR (BL1-A, 448 nm excitation laser, 530/30 nm emission filter) after gating for K562 cells (SSC-A v FSC-A), singlets (FSC-H v FSC-A), Cas9-BFP+ (VL1-A, 405 nm excitation laser, 450/40 emission nm filter), and mCherry-sgRNA+ (YL2-A, 561 nm excitation laser, 620/15 nm emission filter) with an Attune NxT Acoustic Focusing Cytometer. MFI of no doxycycline controls was measured for each genotype.

### Immunoblotting

We employed two methods for immunoblotting. In the first method, K562 cells in RPMI were spun at 800 x g for 5 minutes at 4 °C. Cells were washed with 1 volume PBS and spun for an additional 5 minutes at 800 x g at 4 °C. Pellets were then flash frozen in liquid nitrogen and stored at −80 °C. Cells were lysed in 150 mM NaCl, 20 mM Tris-HCl (7.4 pH), 0.1% NP-40, and Complete^TM^ EDTA protease inhibitor cocktail (Sigma-Aldrich) and incubated shaking at 4 °C for 30 minutes. Lysates were spun at 20,000 x *g* for 15 minutes and supernatants were collected. Samples were normalized using a Pierce 660 nm Protein Assay (Thermo Fisher Scientific) and denatured in 1X NuPAGE LDS sample buffer with reducing agent (Thermo Fisher Scientific) for 10 minutes at 70 °C. Samples were run on 4-15% or 4-20% Mini-PROTEAN SDS-PAGE gels (Bio-Rad, Hercules, CA) in 1X TGS solution (250 mM tris, 192 mM glycine, 0.1% SDS) at 200V for 40-60 minutes. Proteins were transferred onto nitrocellulose membranes for 15 minutes at 1.5A, 25V using a Trans-Blot Turbo Transfer System (Bio-Rad). Membranes were blocked for 1 hour Odyssey® Blocking Buffer (PBS) (LI-COR, Biosciences, Lincoln, NE), and incubated shaking overnight with primary antibodies in 5% BSA in PBST. Membranes were washed three times for 5 minutes with PBST then incubated with 1:10,000 IRDye® 800CW Goat anti-Rabbit IgG (H + L) (LICOR) or IRDye® 800CW Goat anti-Mouse IgG (LI-COR) secondary antibodies in PBST for 1 hour. Membranes were washed three times for 5 minutes with PBST then imaged using an Odyssey CLx Infrared Imaging System (LI-COR). Signal intensities were measured using Image Studio v3 (LI-COR).

In the second immunoblotting method, we adapted a protocol provided by the Lukacs laboratory at McGill University. K562 cells were collected on ice and spun at 800 x *g* for 5 minutes at 4 °C. 1 volume of PBS (with 1 mM CaCl_2_ and 1mM MgCl_2_) was used to wash the cells, and cells were lysed in RIPA buffer (150 mM NaCl, 20 mM Tris, 1% (v/v) Triton-X, 0.1% SDS (w/v), 0.5% (w/v) sodium deoxycholate, pH 7.5) with Complete^TM^ EDTA protease inhibitor cocktail (Sigma-Aldrich). Lysates were incubated on ice for 5 minutes then spun at 20,000 x *g* for 15 minutes at 4 °C. Supernatants were transferred to a new tube, and protein concentrations were normalized using the Pierce BCA Protein Assay Kit (Thermo) per the manufacturer’s instructions. 5X sample buffer (12.5% SDS, 500 mM DTT, 300 mM Tris pH 6.8, 40 mM EDTA, 17.5% glycerol, 3.5% bromophenol blue) was added to the lysates, and the lysates were heated for 15 minutes at 50 °C before being loaded onto 4-15% or 4-20% Mini-PROTEAN SDS-PAGE gels (Bio-Rad, Hercules, CA) in 1X TGS solution (250 mM Tris, 192 mM glycine, 0.1% SDS). Gels were run at 60V for 30 minutes then 130V for 1.5 hours. Proteins were transferred onto nitrocellulose membranes using a Criterion Blotter wet transfer system (BioRad) at 20V overnight or 100V for 2 hours at 4 °C in 25 mM Tris, 192 mM glycine, pH 8.3 with 20% methanol. Membranes were blocked for 1 hour in Odyssey® Blocking Buffer (PBS) (LI-COR) and incubated with primary antibodies in 5% BSA in PBST for 2 hours at 4 °C or RT. Membranes were incubated with secondaries and imaged as detailed above.

Antibodies used in this study were CFTR M3A7 Mouse Monoclonal (Millipore Sigma, MAB3480, 1:1000), CFTR L12B4 Mouse Monoclonal (Millipore Sigma, MAB3484, 1:1000), CFTR 570 Mouse Monoclonal (CFF-UNC CFTR Antibody Distribution Program, A2, 1:1000), CFTR 596 Mouse Monoclonal (CFF-UNC CFTR Antibody Distribution Program, A4,1:1000), GAPDH 14C10 Rabbit Monoclonal (Cell Signaling Technologies, 2118S, 1:5000), RNF5 22B3 Mouse Monoclonal (Santa Cruz Biotechnology, sc-81716, 1:1000), RNF185 + RNF5 Rabbit Monoclonal (Abcam, ab181999, 1:1000), and ɑ-Tubulin Rabbit Polyclonal (Abcam, ab15246, 1:1000-2000),

### Drug studies

For degradation kinetics and steady-state experiments with inhibitors, reporter cells were induced with 1 μg/ml doxycycline for 16 hours then dosed with 20 μM NMS-873 (Sigma-Alrich), 2 μM TAK-243 (Med Chem Express, Monmouth Junction, NJ), or 1 μM bortezomib (BTZ, Selleck Chemicals, Houston, TX) for 3 hours. For experiments with corrector drugs, lumacaftor (VX-809, 2.5 μM), tezacaftor (VX-661, 5 μM, Selleck Chemicals), Corr-4a (2.5 μM, EMD Millipore, Burlington, MA), ivacaftor (VX-770, 1 μM, Selleck Chemicals) and/or elexacaftor (VX-445, 5 μM, Selleck Chemicals) were co-administered with 1 μg/ml doxycycline for 16 hours.

### Lentiviral transduction of the Bassik Lab Human CRISPR-Cas9 Deletion Library into K562 cells

Each of the nine sublibraries comprising the Bassik Lab CRISPR knockout library (Addgene #101926, #101927, #101928, #101929, #101930, #101931, #101932, #101933, #101934) was lentivirally integrated into our K562 reporter cells. For each sublibrary, HEK293T cells were passaged into 15 cm plates at 7.5 x 10^6^ cells per plate in 30 ml DMEM + 10% FBS. The next day, 48 μl of TransIT-LT1 (Mirus Bio LLC) reagent was mixed into 1.3 ml Opti-MEM^TM^ (Thermo Fisher Scientific) and incubated for 5 minutes at room temperature. 8 μg sublibrary plasmids (gift from Michael Bassik) and 8 μg third generation lentiviral packaging plasmid mix (pVSV-G, pMDL, and pRSV at a 1:1:1 ratio) were combined with the Opti-MEM, and the mixture was incubated for an additional 20 minutes at room temperature before being added to the 15 cm plate of HEK293T cells. After 48 hours, the viral supernatant was filtered through a 33 mm 0.45 μM PVDF syringe filter (Millipore Sigma, Cat# SLHVM33RS) and stored at 4 °C or −80 °C. 30 ml of fresh media was added to the cells, and the viral supernatant was collected, filtered, and stored at 4 °C or −80 °C again at 72 hours.

To integrate the virus into the reporter cells, 3.5 x 10^7^ K562 cells expressing Cas9-BFP were resuspended into 60 ml of viral media with 8 μg/ml polybrene (hexadimethrine bromide, Sigma-Aldrich) in 6-well plates. The plates were spun at 1000 x *g* for 2 hours at 33 °C in a M-20 Microplate Swinging Bucket Rotor (Thermo) on a Sorvall Legend XTR Centrifuge (Thermo). The viral media was removed from the cells, and the cells were resuspended into 70 mL of RPMI media + 10% FBS and grown in T-225 cm^2^ flasks. The following day 60 mL of media was added to the cells. On Day 2 after the spin transduction, the MOI was determined by measuring the percent mCherry positive cells using an Attune NxT Acoustic Flow Cytometer using the yellow 561 nm laser with a 620/15 nm filter. If 20-50% of the cells were mCherry positive, 60 mL of cells were removed, and 60 mL fresh media was added. On day 3 and day 5, 60 mL media was removed and 60 mL of media with puromycin (Thermo Fisher Scientific) was added to a final puromycin concentration of 1 μg/mL. After day 6, puromycin media was removed once the cells were > 90% mCherry positive, and cells were expanded from days 7-9. On day 9, the cells were frozen down at 5-10 x 10^6^ cells/mL in 90% FBS, 10% DMSO at −80 °C in 5 ml cryogenic vials, and the next day the vials were transferred to liquid nitrogen for long term storage. Screen replicates were performed with two independent lentiviral integrations per sublibrary.

### Genome-wide CRISPR-Cas9 knockout screens with the Bassik Lab Human CRISPR-Cas9 Deletion Library

Growth media for cells (RPMI media with L-glutamine, no phenol red, 10% FBS, pen-strep) was placed into 37 °C tissue culture incubators 4-12 hours prior to defrosting reporter cells to ensure that the media was pre-gassed with CO_2_, as pre-gassing media was found improve transcriptional shutoff after doxycycline washout. Cryovials per each of the nine sublibraries (∼3-5 x 10^7^ cells) were defrosted in a 37 °C water bath, and cells were spun down for 5 minutes at 800 x *g* to remove the freezing media. Each of the nine sublibraries were individually placed into 100 ml growth media in T-225 cm^2^ flasks, and the concentration of cells was measured using an Attune NxT Acoustic Flow Cytometer. Cells from the 9 sublibraries were combined into 500 mL Micro-Carrier Spinner Flasks (Bellco Glass, Vineland, NJ, Cat#1965-02500) such that each sgRNA library was represented at 1000X (e.g., 2 x 10^7^ cells for a 20,000 sgRNA sublibrary) and the concentration of cells was 3.5 x 10^5^ cells/mL. Cells were spun at 100 rpm using a Bell-ennium Digital Magnetic Stirrer (Bellco Glass, Cat# 7785D2005) at 37 °C in 5% CO_2_. After 24 hours, cells were expanded by diluting to 3.5 x 10^5^ cells/mL in pre-gassed growth media. 48 hours after thawing, cells were spun down at 800 x *g* for 5 minutes in 500 mL polypropylene centrifuge tubes (Corning), and 2 L pre-gassed growth media plus 0.1 ug/mL doxycycline was added to a final cell concentration of 3.5 x 10^5^ cells/mL to ensure ∼4000X coverage of the genome-wide library at the start of the sorting experiment, and the cell suspension was split between four 500 mL spinner flasks. After 12 hours of doxycycline induction, cells were spun down at 800 x *g* for 5 minutes at 37 °C and cells were washed with pre-gassed 500 ml RPMI media, with L-glutamine, no phenol red (without FBS and Pen-Strep). This wash step was repeated for a total of two washes, and the cells were placed in 2 L growth media without doxycycline in spinner flasks. The doxycycline washout was monitored every hour for 4 hours using an Attune NxT Acoustic Flow Cytometer, and after 4.5 hours, the cells were spun down at 800 x *g* for 5 minutes to remove the growth media and placed on ice to halt degradation.

Cells were resuspended ice-cold RPMI media, with L-glutamine, no phenol red with 0.5% FBS to a final concentration of 1-1.5 x 10^7^ cells/mL, and the resuspension was filtered through 70 μM cell strainers and placed on ice. Cells were sorted on two BD FACSAria II Cell Sorters (BD Biosciences) equipped with a blue 488 nm laser with a 505 nm long pass mirror and 525/50 nm filter (for mNG), a green/yellow 532 nm laser with a 600 long pass mirror and 610/20 nm filter (for mCherry), and a violet 405 nm laser with a 450/50 nm filter (for BFP). The following gating hierarchy was used to sort for the top 5% mNG populations: SSC-A v FSC-A polygonal gate for live K562 cells, FSC-H v FSC-A polygonal gate for singlets, V450-A histogram gate for BFP+ cells, G610-A histogram gate for mCherry + cells, and B525-A histogram gate for the top 5% mNG+. Samples were kept at 4 °C and agitated at 300 rpm inside the sorters. Cells were sorted into 15 ml conical tubes with 1-5 ml of RPMI, L-glutamine, no phenol red + 30% FBS. Sorting was performed using an 85 μm nozzle at 10-15,000 events/second with 4-way purity. Approximately ∼1.2 x 10^6^ cells were collected during sorting to ensure 1000X coverage of the top 5% of the ∼220,000 element genome-wide library, and ∼2-4 x 10^8^ unsorted cells were collected to ensure 1000X-2000X coverage of the full library.

After sorting, cells were spun down at 3,000 x *g* for 15 minutes at 4 °C then washed with ice-cold 5 mL PBS. The cells were spun again at 3,000 x *g* for 15 minutes at 4 °C. After the PBS was removed, cells were transferred to 1.5 mL microcentrifuge tube and the excess PBS was removed through an additional spin at 3,000 x *g* for 15 minutes at 4 °C. Cells were frozen with liquid nitrogen and stored at −80 °C.

### Preparation of NGS sequencing libraries for Bassik Lab Human CRISPR-Cas9 Deletion Library

For sorted populations with < 5 x 10^6^ cells, genomic DNA was isolated using QIAamp DNA mini kits (Qiagen, Hilden, Germany) according to the manufacturer’s protocol, eluting in 200 μl EB buffer (10 mM Tris-Cl, pH 8.5) instead of buffer AE. For sorted populations with 0.5-2 x 10^7^ cells, genomic DNA was isolated using QIAamp DNA midi kits (Qiagen) according to the manufacturer’s protocol with the following modifications: 1) QIAmp midi columns were spun at 4000 rpm for 3 minutes after the addition of buffer AW1, 2) columns were spun at 4000 rpm for 25 minutes after the addition of buffer AW2, 3) columns were incubated with 200 μl EB buffer for 5 minutes then spun at 4000 rpm for 5 minutes, and 4) columns were incubated with an additional 150 μl EB buffer for 5 minutes then spun at 4000 rpm for 5 minutes.

For unsorted populations (0.2-4 x 10^8^ cells), genomic DNA was isolated using QIAamp DNA maxi kit components (Qiagen). Unsorted cells were divided such that each column purified genomic DNA from no more than 1 x 10^8^ cells. For each purification, cells were resuspended in 6.25 ml room temperature PBS. 500 μl QIAGEN protease and 6 ml buffer AL were added and samples were shaken vigorously by hand for 2 minutes. Samples were incubated in a 70 °C water bath for 10 minutes before 5 ml 100% ethanol was added, the sample inverted 10X immediately after the addition of ethanol. The mixture was loaded onto a QIAamp Maxi spin column and spun at 3900 rpm for 3 minutes. The flow through was removed, and 5 ml of buffer AW1 was added to the column and the column was spun at 3900 rpm for 2 minutes. After the spin, the flow through was discarded and 5 ml buffer AW2 was added, and the column was spun at 3900 rpm for 20 minutes. The column was transferred to a new 50 ml conical tube, and the columns were incubated for 5 minutes with 800 μl buffer EB then spun at 3900 for 5 minutes. The elution step was repeated with an additional 800 μl buffer EB, and DNA purifications from the same population were combined. The DNA concentration, A260/280 ratio, and A230/280 ratio for each sample was measured using a NanoDrop 2000 Spectrophotometer (Thermo), and samples were diluted to 250 ng/ul.

Nested PCR was used to amplify the sgRNAs from the genomic DNA. To ensure the highest coverage in sequencing, all genomic DNA harvested during the sorting was amplified during the PCR1 step with oMCB1562 (AGGCTTGGATTTCTATAACTTCGTATAGCATACATTATAC) and oMCB1563 (ACATGCATGGCGGTAATACGGTTATC). Genomic DNA was normalized to 250 ng/µl, and 40 µl (10 µg) was added to each 100 µl PCR1 reaction. If the concentration was less than 250 ng/µl, 40 µl of the gDNA was added to each reaction. For each reaction, 20 µl 5X Herculase Buffer (Agilent, Santa Clara, CA), 1 µl 10 mM dNTPs (with the concentration of each nucleotide at 10 mM), 1 µl 100 µM oMCB1562, 1 µl 100 µM oMCB1563, 2 µl Herc II polymerase (Agilent), and 35 µl nuclease-free water were added to the 40 µl gDNA, and the reaction was amplified 1 x 98 °C/2 min; 18 x 98 °C/30 s, 59.1 °C/30 s, 72 °C/45 s; and 1 x 72 °C/3 min. PCR1 reactions were combined into a single tube, and 5 µl of PCR1 was added to the PCR2 reaction. To each 100 µl PCR2 reaction, 20 µl 5X Herculase Buffer, 2 µl 10 mM dNTPs, 0.8 µl 100 µM oMCB1439 (CAAGCAGAAGACGGCATACGAGATGCACAAAAGGAAACTCACCCT), 2 µl Herc II polymerase, 0.8 µl 100 µM barcoded CRISPR KO primer (oMCB1440, aatgatacggcgaccaccgagatctacacGATCGGAAGAGCACAC GTCTGAACTCCAGTCAC NNNNNN CGACTCGGTGCCACTTTTTC, where NNNNNN is an Illumina TruSeq index), and 69.4 µl nuclease-free water were added to the 5 µl PCR1 reaction, and the reaction was amplified 1 x 98 °C/2 min; 19 x 98 °C/30 s, 59.1 °C/30 s, 72 °C/45 s; and 1 x 72 °C/3 min. PCR2 products were run on a 2% TBE agarose gel at 150 V for 1 hour, and the 280 bp PCR2 product was purified using the QIAquick Gel Extraction Protocol (Qiagen), according to the manufacturer’s protocol with the following modifications: 1) 15 µl 3 M sodium acetate added to the buffer QC before adding isopropanol, and 2) the samples were eluted in 50 µl buffer EB. Samples were quantified using a Qubit 2.0 or 3.0 (Thermo) using the Qubit HS dsDNA Assay (Thermo). Samples were pooled and normalized with water into a 1 nM library.

PCR amplicons were sequenced using a NextSeq 550 (Illumina, San Diego, CA) using a NextSeq 500/550 High Output Kit v2.5 (75 Cycles). To prepare the library for sequencing, 20 µl 1 nM library was added to 20 µl freshly diluted 0.2 M NaOH then vortexed, spun at 250 x g for 1 min, and incubated at room temperature for 5 minutes. 20 µl 0.2 M Tris-HCl, pH 7 was added, and the sample was vortexed and spun down at 250 x g for 1 min. 940 µl ice-cold buffer HT1 (Illumina) was added to generate a 20 pM library, and the sample was inverted to mix. To generate the 2.2 pM library (a higher concentration than the recommended 1.8 pM), 130 µl library was added to 1170 µl Buffer HT1. 6 µl of µM100 custom, PAGE-purified sequencing primer, oMCB1672 (GCCACTTTTTCAAGTTGATAACGGACTAGCCTTATT TAAACTTGCTATGCTGTTTCCAGCTTAGCTCTTAAAC), was diluted in 2 mL Buffer HT1 to a final concentration of 0.3 µM. The samples were sequenced using 21 nt single reads and 6 nt index reads, with custom primers for the single reads. Only sequencing runs with a cluster density of 100-500 K/mm^2^ and % cluster passing filter > 80% were processed for downstream analysis. Sequencing runs were processed into fastq files by index using bcl2fastq (Illumina).

### sgRNA enrichment analysis with casTLE

casTLE sgRNA enrichment analysis (Morgens *et al*., 2016) was performed using a SLURM-based HPC-cluster CRISPR/Cas9 screening analysis pipeline (Kramer *et al*., 2018; Kim *et al*., 2022) on Sherlock, a high-throughput computing cluster managed by the Stanford Research Computing Center. The casTLE maximum likelihood estimation function was run to generate p-values, using 10^6^ and 10^5^ permutations for the genome-wide and UBAL screens, respectively. Adjusted p-values (FDR) were calculated using the Benjamini-Hochberg procedure in R. Custom lists of E1, E2, and E3 genes were compiled for downstream bioinformatics analysis in R and for generating the UBAL sublibrary. A list of GO:0006402 mRNA catabolic process genes was acquired from BioMart, and a curated list of chaperone/co-chaperone genes was downloaded from (Shemesh *et al*., 2021).

### qPCR analysis of CFTR transcript abundance

After mNG-F508del was induced with doxycycline for 16 hours, approximately 5-10 x 10^5^ K562 cells were spun at 800 x g for 5 minutes and lysed in TRIzol reagent (Invitrogen). mRNA was purified using the Direct-zol™ RNA Miniprep Plus Kit (Zymo) according to the manufacturer’s instructions. cDNA synthesis and RT-qPCR was carried out using Luna® Universal One-Step RT-qPCR Kit (NEB) according to the manufacturer’s instructions with 100 ng of mRNA. Relative fold changes were calculated in technical replicates using the 2^-ΔΔCt^ method with *GAPDH* as the reference gene. *CFTR* qPCR forward and reverse primers were TCTCCTTTCCAACAACCTGAACAAA and CTCCCAGATTAGCCCCATGAG (Masvidal *et al*., 2014) while the GAPDH pPCR primers were TGTCGCTGTTGAAGTCAGAGGAGA and AGAACATCATCCCTGCCTCTACTG.

### Creation of pooled single and double knockout cell lines using Cas9 RNP electroporation

Pooled knockout cell lines were generated by electroporating Cas9-guide RNA ribonucleoprotein complexes into reporter K562 cells using an Amaxa 4D Nucleofector X Unit (Lonza) and the SF Cell Line 4D-Nucleofector X Kit S (Lonza). Gene Knockout Kits v2 (Synthego, Redwood City, CA) composed of 3 unique guide RNAs per target were ordered for each gene of interest. Guide RNAs were resuspended in 10 mM Tris, 1 mM EDTA pH 8.0 to a final concentration of 100 µM. Cas9 RNPs were generated by incubating 0.6 µL SpyFi Cas9 (Aldevron, South Fargo, ND) and 1 µL (3.2 µg) guide RNA per gene target at room temperature for 10 minutes prior to electroporation. 2 x 10^5^ cells were resuspended in 20 µL Nuclofector Solution SF (Lonza), combined with the assembled Cas9 RNPs, and the cells were electroporated in a 16-well cuvette using program FF-120.

Variable guide RNA sequences included in the Synthego Kits for *RNF185*, *RNF5*, and *UBE2D3* were: CAGCCAAGGAUGGCAAGCAA (RNF185 #1), AAUGGCGCUGGCGAGAGCGG (RNF185 #2), CAGGCUGAUGACGGCAUCCU (RNF185 #3), GUCUCUCACCUGGGAUCCUG (RNF5 #1), UCUUCCACACCGUUUUCCAA (RNF5 #2), GGCUGGAGACACGGCCAGAA (RNF5 #3), UAGAGCAUUCUUGGAAGAUA (UBE2D3 #1), UGAGGGAAAAUACUUGCCUU (UBE2D3 #2), and CAGAAUGACAGCCCAUAUCA (UBE2D3 #3). A synthetic sgRNA against the *AAVS1* safe-harbor locus served as a control guide (guide sequence: GGGGCCACUAGGGACAGGAU ((Amrani *et al*., 2018)).

### Creation of pooled single knockout cell lines using lentivirus

sgRNAs of interest were cloned into pMCB320 (Addgene #89359, (Han *et al*., 2017)), which serves as the backbone for the Bassik Human CRISPR/Cas9 Deletion plasmid library. pMCB320 expresses a tracrRNA from a mouse U6 promoter and contains an mCherry-T2A-Puro selection marker. To clone a sgRNA into this backbone, “top” and “bottom” oligos were designed with BstXI and modified stem loop/BlpI overhangs. The sequence of the top oligo was 5’-TTG<u>G</u>-(variable sgRNA sequence)-GTTTAAGAGC-3’ while the sequence of the bottom oligo was 5’-TTAGCTCTTAAAC-(reverse complement of the variable sgRNA sequence)-<u>C</u>CAACAAG-3’, with the underlined “G” representing the first G of the variable sgRNA sequences listed in the Bassik Human CRISPR/Cas9 Deletion Library. To phosphorylate the oligos, 1 µL100 µM top oligo, 1 µL 100 µM bottom oligo, 1 µL T4 DNA ligation buffer (NEB) 6.5 µL water, and 0.5 µL T4 PNK (NEB) were combined and incubated at 37 °C for 30 minutes followed by 95 °C for 5 minutes in a thermocycler. The oligos were subsequently annealed by ramping down the temperature at 0.1 °C per second 700X to 12 °C.

pMCB320 was digested with FastDigest BstX1 and BlpI (Thermo Fisher Scientific) for 15 minutes at 37 °C then dephosphorylated with Fast AP for 15 minutes at 37 °C (Thermo Fisher Scientific). The digested vector was gel-purified using QIAquick Gel Extraction Kit (Qiagen), and 1 µL of a 1:500 dilution of the annealed oligos were added to 50 ng digested vector, T4 DNA ligase (NEB), and T4 DNA ligase buffer (NEB) in a 11 µL reaction and ligated for 1 hour at 25 °C followed by 10 minutes at 65 °C. K562 cells were transduced with lentivirus generated from pMCB320 using the “Small-Scale Lentiviral Transduction of K562 Cells” protocol detailed above, and K562 cells were incubated with 1 µg/mL puromycin for 6-9 days until > 90% mCherry positive.

The variable sgRNA sequences from the Bassik Human CRISPR/Cas9 Deletion Library used for our study were GAAGCAAAACTTCAGACTAC (sgSAFE.6665, 0Safe_safe_UNA3_211214.6665), GAAGAAAAGTCTATTCATAT (sgUBA5.3, ENSG00000081307_UBA5_PROT_80782.3), GTGCAATCTCTGGCACTAGG (sgUBE2D3.2, ENSG00000109332_UBE2D3_PROT_71923.2), and GCTCTCCGGAGCTGGTCTC (sgRNF5.3, ENSG00000204308_RNF5_PROT_167655.3).

### Creation of pooled double-knockout cell lines using lentivirus

Double-knockout cell lines were generated as detailed previously (Han *et al*., 2017). One sgRNA was cloned into pMCB320 as detailed above (see “Generation of Single Knockout Cell Lines with sgRNAs from the Bassik Human CRISPR/Cas9 Deletion Library”) while a second sgRNA was cloned into pKHH030 (Addgene #89358). To clone a sgRNA into pKHH030, “top” and “bottom” oligos were designed with BbsI sites. The sequence of the top oligo was 5’-ACC<u>G</u>-(variable sgRNA sequence)-3’ while the sequence of the bottom oligo was 5’-AAAC-(reverse complement of the variable sgRNA sequence)-3’, with the underlined “G” representing the first G of the sgRNA sequences listed in the Bassik Human CRISPR/Cas9 Deletion Library. Oligos were phosphorylated and annealed as detailed above. 1 µg pKHH030 was digested with BbsI (NEB) in CutSmart Buffer (NEB) in a 50 µl reaction for 15 minutes at 37 °C then dephosphorylated with 1 µl Fast AP for 15 minutes at 37 °C before heat inactivation for 5 minutes at 75 °C. The digested vector was gel-purified using QIAquick Gel Extraction Kit (Qiagen), and 1 µL of a 1:250 dilution of the annealed oligos were added to 50 ng digested vector, T4 DNA ligase (NEB), and T4 DNA ligase buffer (NEB) in a 11 µL reaction and ligated for 1 hour at 25 °C followed by 10 minutes at 65 °C before transformation into DH5 alpha bacteria.

pKHH030 and pMCB320 carrying the desired sgRNA sequences were double digested with XhoI (NEB) and BamHI-HF (NEB) in CutSmart Buffer at 37 °C for 5-15 minutes followed by heat-inactivation at 65 °C for 20 minutes. The insert from pKHH030 and the vector from pMCB320 were gel purified then ligated with T4 DNA ligase at 16 °C overnight followed by heat inactivation at 10 minutes at 65 °C. L. Transduction and puromycin selection of double-knockout K562 cells was conducted as detailed above for single knockout K562 cells.

### Creation of pooled double-knockout cell lines for sensitized screens

Dual guide pMCB320 plasmids were transiently expressed in mNG-F508del Cas9-BFP K562 cell lines, and BFP+/mCherry+ cells were sorted to generate stable pools of knockout cell lines. sgSAFE.6665/sgSAFE.6665, sgRNF5.3/sgSAFE.6665, sgUBE2D3.2/sgSAFE.6665, and sgRNF5.3/sgUBE2D3.2 dual guide RNA plasmids were generated using the pMCB320 backbone as detailed above, and K562 cells were electroporated with a Nucleofector 2b device (Lonza, Basel, Switzerland) using the Cell Line Nucleofector^TM^ Kit V (Lonza) according to the manufacturer’s protocol for K562 cells. 1 x 10^6^ cells were centrifuged at 800 x *g* for 5 minutes, resuspended in 100 µL Cell Line Nucleofector Solution V, combined with 2 µg plasmid DNA, and electroporated using program T-016. Cells were resuspended in 500 µL RPMI, and after 24 hours, the cells were prepared for sorting as detailed above. Cells were sorted on a BD FACSAria II using the same protocol for the genome-wide screens except the 525 nm channel was not activated because mNG was not induced with dox.

### Construction of <u>u</u>biquitome/<u>a</u>utophagy/lysosomal (UBAL) degradation sgRNA library

The custom UBAL degradation library (**Table S2**) contains 20,710 elements with 18,710 sgRNAs targeting 1,871 genes (∼10 sgRNAs per gene) and 2,000 negative control sgRNAs selected from the Bassik Lab Human CRISPR-Cas9 Deletion Library. The custom library was constructed as previously described (Morgens *et al*., 2016, 2017). Synthesized oligonucleotides (Twist Biosciences, South San Francisco, CA) were PCR-amplified for 10 cycles using KAPA HiFi HotStart DNA Polymerase (Roche, Basel, Switzerland) with 52 °C annealing and 15 s extension. PCR products were purified with a Qiagen MinElute PCR Purification Kit (Qiagen), and eluted samples were digested with BstXI and BlpI restriction enzymes overnight at 37°C. Digests were run on a 20% TBE gel (Thermo Fisher Scientific). The 33 bp band was excised, purified over a Co-star Spin X column (Corning), precipitated using isopropanol, resuspended in Qiagen Buffer EB, and ligated into BstXI/BlpI-cut pMCB320 using T4 DNA ligase. Ligations were transformed into Endura Electrocompetent Cells (LCG Biosearch Technologies, Middlesex, UK) using a Gene Pulser II (Bio-Rad) set at 1.8 kV, 600 U, 10 mF. Cells were recovered for 1.5 h at 37 °C, plated on 500-cm^2^ agar plates with 75 mg/ml carbenicillin, and grown overnight at 37 °C. Colonies were scraped off plates, and library plasmids were purified using a Qiagen HiSpeed Maxi kit (Qiagen, 12662). Plasmids were eluted in Buffer EB, aliquoted, and stored at −80 °C.

### Sensitized screens with RNF5^KO^, UBE2D3^KO^, and RNF5/UBE2D3^KO^ reporter cell lines

Pooled knockout cells generated using transient transfection of dual-guide RNA plasmids were transduced with the UBAL sublibrary using the same protocols for integrating the Bassik Lab CRISPR-Cas9 Human Deletion Library (see above). The sensitized screens with the four genotypes were conducted in tandem in two separate experiments using the same protocols for the genome-wide screens with the following modifications: 1) reporter shutoff was initiated by dosing the cells with 20 µM emetine for 90 minutes, and 2) mCherry+ cells were sorted using a blue 488 nm laser with 595 nm long pass mirror and 610/20 nm filter. Approximately ∼1 x 10^5^ cells were collected during sorting to ensure 1000X coverage of the top 5% of the ∼2,000 element genome-wide library, and ∼1-2 x 10^7^ unsorted cells were collected to ensure 1000X-2000X coverage of the UBAL sublibrary.

## ACKNOWLEDGEMENTS

We would like to thank the Stanford Shared FACS Facility, and the Stanford Research Computing Center for providing core resources for this research. We also acknowledge Josh Tycko, David Morgens, Kaitlyn Spees (Bassik laboratory), Garam Kim, Julien Couthouis (Gitler laboratory), Laura Rojas (Milla laboratory), Francesco Scavone, Sam Gumbin, Paul Da Rosa (Kopito laboratory), Michael Haney (Wyss-Coray laboratory), Melissa Roberts (Olzmann laboratory), and Leslie Magtanong (Dixon laboratory) for help with this project. This work was supported by 5R01-GM074874 (RRK), 2R56-NS042842 (RRK), and R01DK128099 (JAO). Celeste Riepe was funded by the Cystic Fibrosis Foundation Path-to-a-Cure Postdoctoral Fellowship.

Anais Amaya was supported by the Elizabeth Nash Memorial Fellowship from the Cystic Fibrosis Research Institute and Postdoctoral Support from the Stanford Maternal and Child Health Research Institute. Operation of the Next Seq 550 was supported through a Bertarelli foundation gift to Stanford’s Brain Rejuvenation Project managed through the Weiss-Coray and Gitler laboratories. The BD FACSAria II sorters at the Stanford Shared FACS facility were supported by NIH Shared Instrumentation grant S10RR025518-01.

## SUPPLEMENTAL INFORMATION

Table S1: casTLE Analysis of genome-wide CRISPR knockout screens with mNG-F508del

**Sheet 1:** casTLE analysis comparing the sgRNA frequencies in the top 5% mNG-F508del positive cells with those in the unsorted populations from two replicate genome-wide CRISPR knockout screens.

Sheet 2: Significant genes from genome-wide screen (FDR < 1%). Sheet 3: Sig. E1 ubiquitin activating enzyme genes. Sheet 4: Sig. E2 ubiquitin conjugating enzyme genes. Sheet 5: Sig. E3 ubiquitin ligase genes. Sheet 6: Sig. chaperone/co-chaperone genes. Sheet 7: Sig. mRNA decay genes.

Table S2: Ubiquitome, autophagy, and lysosome (UBAL) degradation sgRNA sublibrary

**Sheet 1**: Legend

**Sheet 2:** List of 1,871 genes in the library (∼10 guides per gene)

**Sheet 3:** Sequences for 20,710 elements (including 2,000 negative control sgRNAs)

Table S3: casTLE Analysis of UBAL sublibrary screens

**Sheet 1:** casTLE analysis comparing the sgRNA frequencies in the top 5% mNG-F508del positive cells with those in the unsorted populations from two replicate control UBAL screens. **Sheet 2:** RNF5-KO UBAL screen analysis. **Sheet 3:** UBE2D3-KO UBAL screen analysis. **Sheet 4:** RNF5/UBE2D3-KO UBAL screen analysis. **Sheet 5:** Significant genes identified in control screens (FDR < 1%). **Sheet 6:** Sig. genes for RNF5-KO. **Sheet 7:** Sig. genes for UBE2D3-KO. **Sheet 8:** Sig. genes for RNF5/UBE2D3-KO screens.

**Figure S1:**
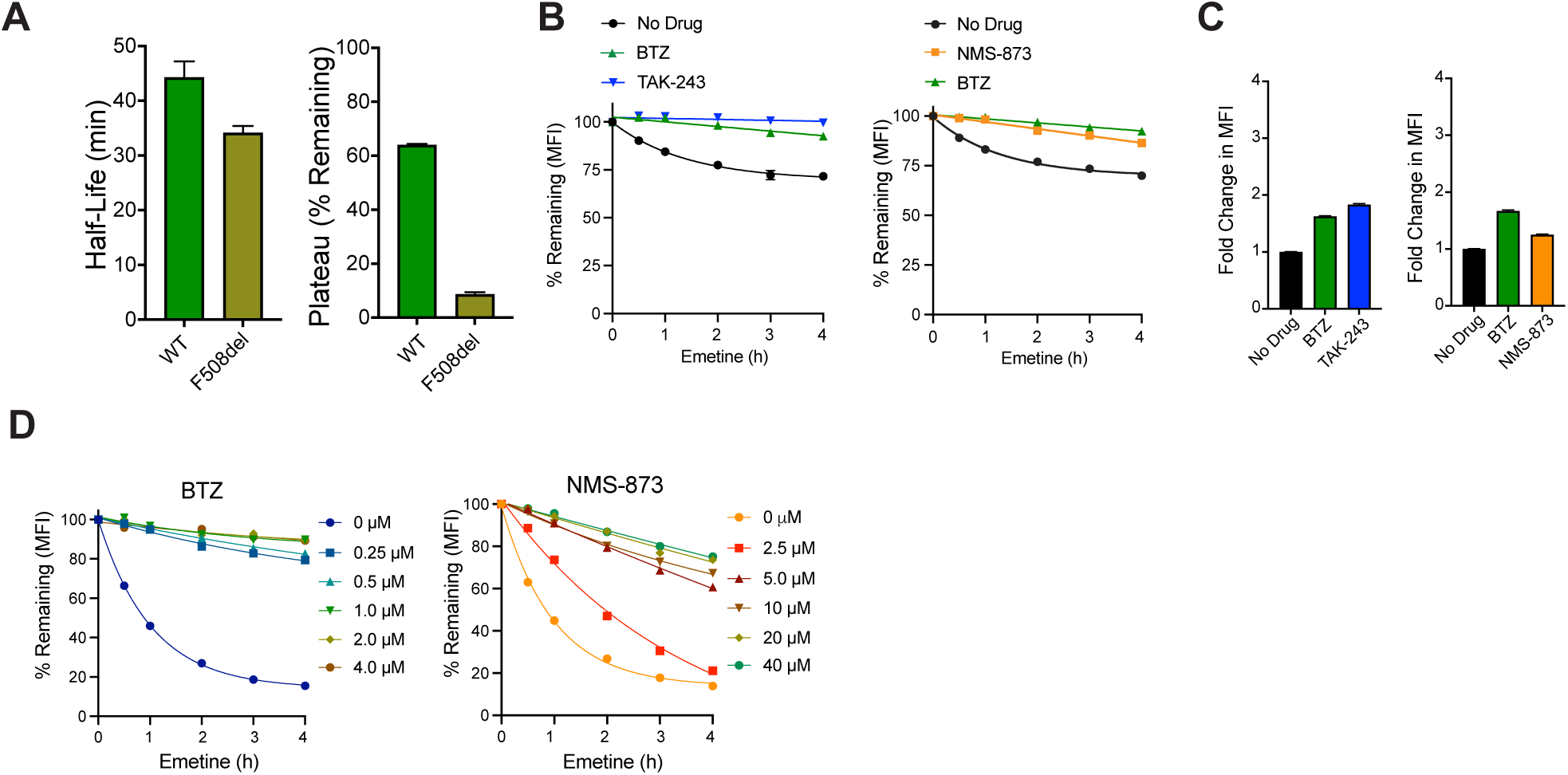
Supplementary information for. Figure 1. (**A**) Quantification of mNG-WT and mNG-F508del half-life and plateau data from Figure 1D using a one-phase decay model. One curve was drawn for each biological replicate (n = 3) to calculate the half-life and plateau. (**B**) Effect of TAK-243, BTZ, and NMS-873 on mNG-WT degradation kinetics. As in Figure 1E, kinetics of mNG fluorescence decay following translation shutoff. Fold change in MFI after 3 hours of proteasome inhibition with 1 μM bortezomib (BTZ) (left and right), E1 inhibition with 2 μM TAK-243 (left), and VCP inhibition with 20 μM NMS-873 (right). (**C**) Effect of TAK-243, BTZ, and NMS-873 on steady-state levels of mNG-WT. As in Figure 1F, measurements taken at t = 0 prior to the addition of emetine. (**D**) Effect of NMS-873 and BTZ concentration on mNG-F508del degradation. As in Figure 1E, translational shutoff assay of mNG-F508del after administration of NMS-873 and BTZ for 3 hours at different inhibitor concentrations (n = 1 biological replicate, curve = one-phase decay).

**Figure S2:**
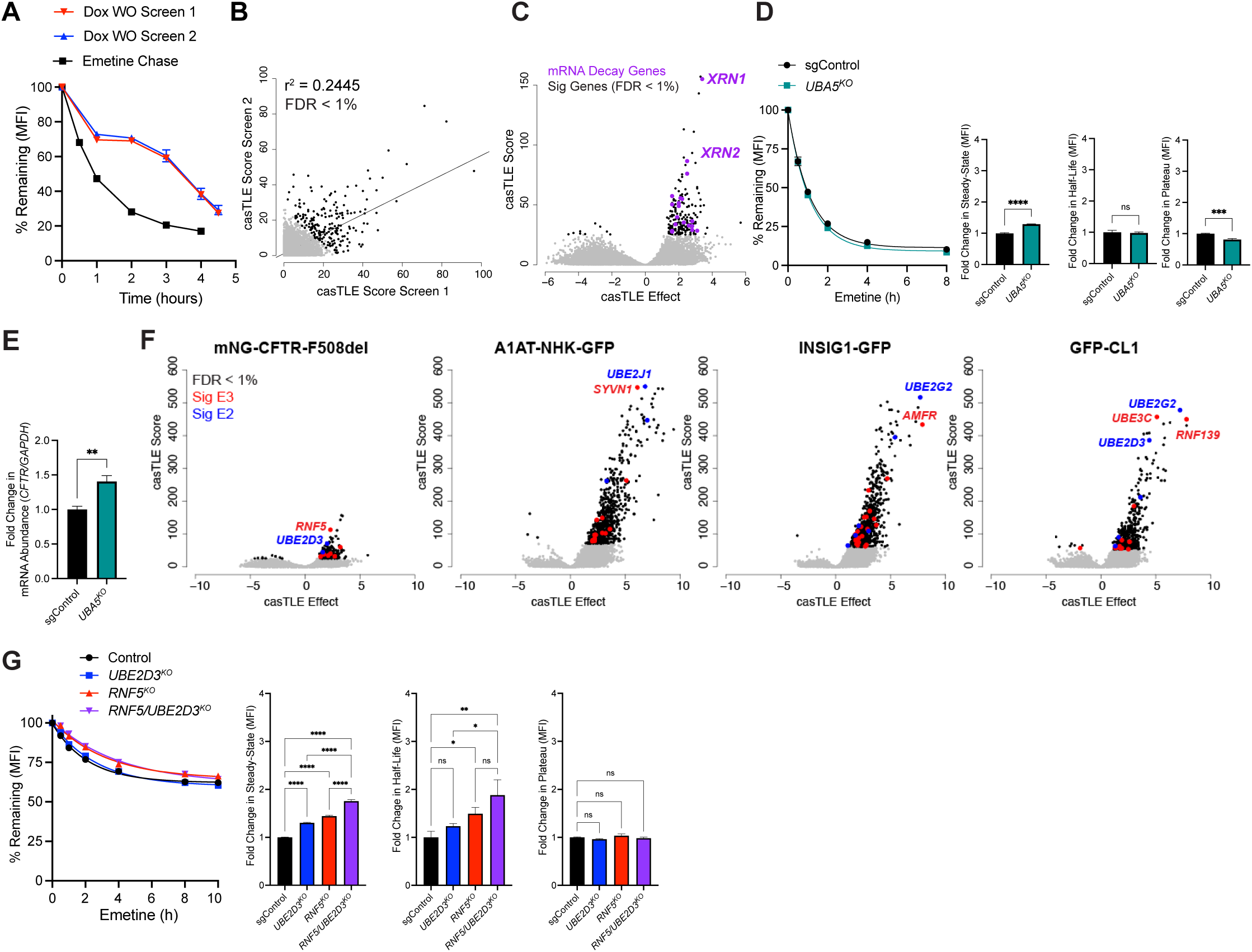
Supplementary information for. Figure 2. (**A**) Doxycycline washout curve for genome-wide screens overlaid with a translational shutoff curve. Translational inhibition with emetine and dox washouts were conducted 12 hours post induction with 0.1 μg/ml dox. (n = 4 spinner flasks per doxycycline washout screen, error bars = SD, n = 1 for translational shutoff). (**B**) Correlation between casTLE scores from screen replicates. Curve = linear regression. Grey = genes observed in screens. Black = significant genes (FDR < 1%). (**C**) Volcano plot of casTLE analysis of single knockout CRISPR screens. Data as in Figure 2B. Purple = significant mRNA catabolic process (GO:0006402) genes. Grey = genes observed in screens. Black = significant genes (FDR < 1%). (**D**) Knocking out the gene encoding the UFMylation activating enzyme, *UBA5*, does not increase mNG-F508del stability. Pooled *UBA5^KO^* reporter cell lines generated using lentiviral integration and puromycin selection of guide RNA transgenes in mNG-F508del reporter cells expressing Cas9-BFP. sgControl = a safe-targeting negative control guide RNA selected from the Bassik Lab Human CRISPR-Cas9 Deletion Library (sgSAFE.6665). Knockout efficiency of sgUBA5 validated in (Walczak et al., 2019). Methods and quantification are as in Figure 2C-D. (*** = p < 0.001, **** = p < 0.0001, unpaired t-test). (**E**) Knocking out *UBA5* increases mNG-F508del transcript abundance. qPCR analysis of *UBA5^KO^* reporter cell lines. Relative fold changes were calculated using the 2^-ΔΔCt^ method with *GAPDH* as the reference gene (n = 3 technical replicates, **** = p < 0.0001, unpaired t-test). (**F**) The casTLE gene effects and scores are lower for mNG-F508del CRISPR knockout screens than those of model ERAD substrates. Data replotted from (Leto et al., 2019). Grey = genes observed in screens. Black = significant genes (FDR < 1%). Red = significant E3 ligases. Blue = significant E2 conjugating enzymes. (**G**) Knocking out *RNF5* and *UBE2D3* singly or in combination have modest effects on mNG-WT degradation. Pooled single and double knockout mNG-WT Cas9-BFP reporter cell lines generated using lentiviral integration and puromycin selection of dual-guide RNA transgenes. sgControl = a dual guide construct expressing two safe-targeting negative control guide RNAs (sgSAFE.6665). Degradation kinetics methods and quantification are the same as in Figure 2C-D.

**Figure S3:**
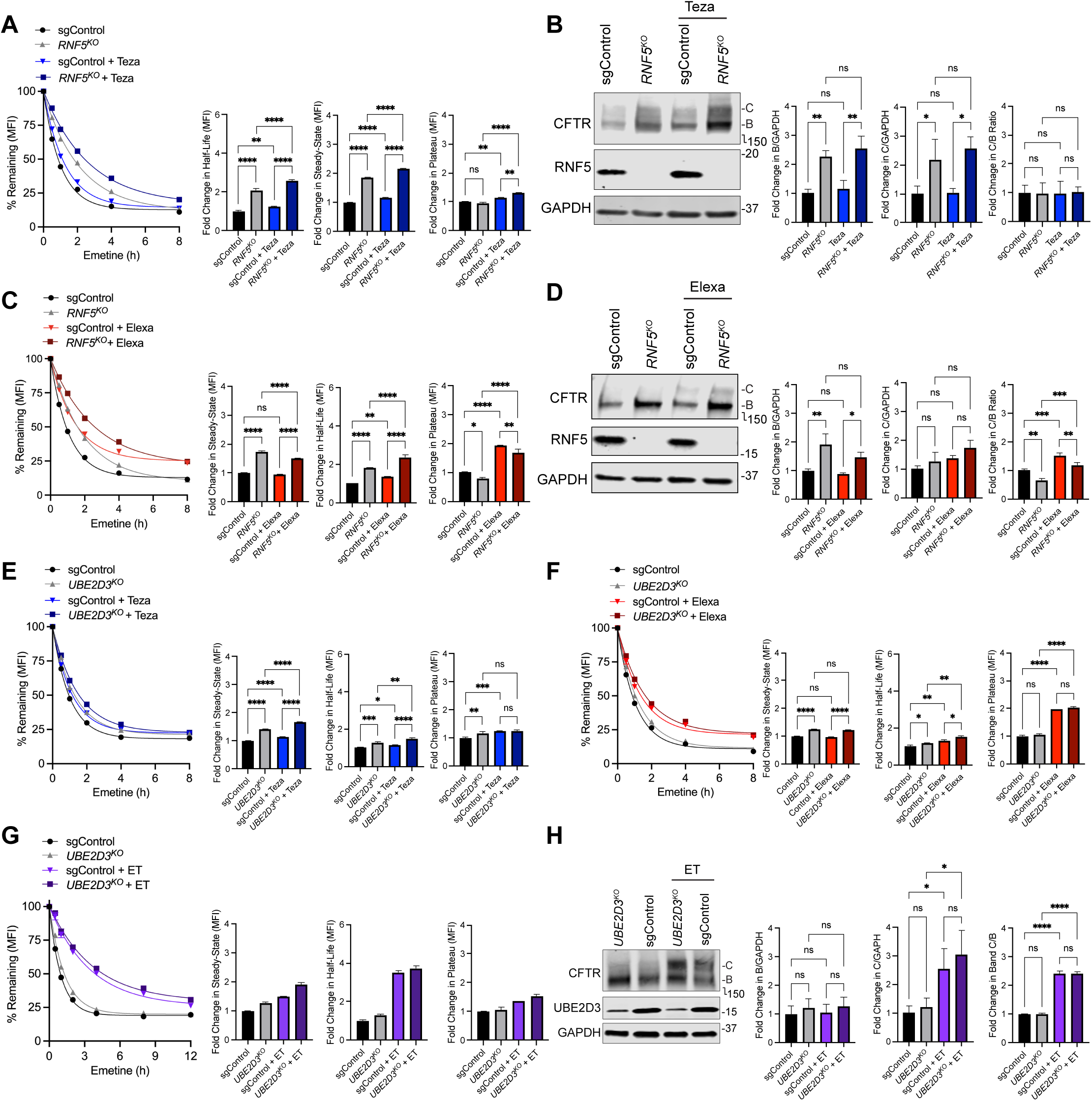
Supplemental data for. Figure 4. (**A-H**) Knocking out *RNF5* or *UBE2D3* has an additive effect with tezacaftor and elexacaftor on mNG-F508del half-life (**A**, **C**, **E**, **F**, **G**) and maturation (**B**, **D**, **H**). Cells were administered tezacaftor (VX-661, 5 μM) and/or elexacaftor (VX-445, 5 μM) for 16 hours. Methods and quantification are as in Figure 2C-F.

